# Mechanical intercellular communication via matrix-borne cell force transmission during vascular network formation

**DOI:** 10.1101/2021.08.17.456669

**Authors:** Christopher D. Davidson, Samuel J. DePalma, William Y. Wang, Jordan L. Kamen, Danica Kristen P. Jayco, Brendon M. Baker

## Abstract

Intercellular communication is critical to the development and homeostatic function of all tissues. Previous work has shown that cells can communicate mechanically via transmission of cell-generated forces through their surrounding extracellular matrix, but this process is not well understood. Here, we utilized synthetic, electrospun fibrous matrices in conjunction with a microfabrication-based cell patterning approach to examine mechanical intercellular communication (MIC) between endothelial cells (ECs) during the assembly of microvascular networks. We found that cell force-mediated matrix displacements in deformable fibrous matrices underly directional migration of neighboring ECs towards each other prior to the formation of stable cell-cell connections. We also identified a critical role for intracellular calcium signaling mediated by focal adhesion kinase and TRPV4 during MIC that extends to multicellular assembly of vessel-like networks in 3D fibrin hydrogels. The results presented here are critical to the design of biomaterials that support cellular self-assembly for tissue engineering applications.

## INTRODUCTION

The ability of cells to communicate and coordinate their activity is crucial to the development and homeostatic function of all tissues (*1*). Intercellular communication through receptor-ligand engagement at the cell-cell interface or via diffusive soluble factors has been extensively studied (*2–5*). In addition to these well-established means of biochemically mediated intercellular signaling, a more recent body of evidence has shown that cells can also communicate via cell-generated forces transmitted to neighboring cells through the extracellular matrix (ECM) (*6, 7*). Cells mechanically engage their surrounding matrix through integrin-based adhesion complexes, or focal adhesions (FAs), which connect the ECM to the cell’s actomyosin cytoskeleton (*8*). These mechanochemical signaling hubs allow cells to continuously sense both passive mechanical and topographical properties of the matrix as well as active external forces applied to the cell (*9*). Concurrently, cell-generated forces applied to the ECM through FAs result in matrix deformations that may impact surrounding cells. The dynamic and reciprocal nature of generating and sensing mechanical signals, however, makes mechanical intercellular communication (MIC) difficult to investigate.

Several prior studies suggest that cell force-generated deformations of the ECM mediate communication between neighboring cells to regulate critical cell functions in cell migration and multicellular assembly. MIC has been observed in a variety of settings spanning different cell types, distinct ECM settings, and across scales ranging from tissues (*10, 11*), to multicellular clusters (*12, 13*), to single cells (*14–18*). For example, in seminal work nearly four decades ago, Stopak and Harris observed that cultured contractile tissue explants embedded within collagen matrices physically reorganize and align collagen fibrils, generating tensile regions that direct cell migration between adjacent explants over millimeter length scales (*10*). At the single cell level, Reinhart-King et al. showed evidence that endothelial cell (EC) traction forces create local gradients of tension that influence EC migration and the formation of contacts between neighboring cells on compliant polyacrylamide gels (*14*). Despite this breadth of evidence, however, we lack an understanding of the cellular machinery required for cells to sense and respond to mechanical signals. Further, how tissue-relevant matrix properties mediate the transmission of cell-generated forces has not been established.

Several computational models suggest that fibrous matrices are optimal for transmitting forces over large distances (i.e., greater than one cell body away) due to their nonlinear elastic behavior and the potential for strain-induced alignment of ECM fibers (*19–28*). Indeed, collagen and fibrin hydrogels, materials that possess fibrous microstructure and nonlinear mechanics, are the primary settings where long-range mechanical interactions between cells have been previously documented. However, the limited structural and mechanical tunability of these materials limits understanding of how ECM properties regulate MIC (*29*). Conversely, tunable, non-fibrous synthetic hydrogels such as polyacrylamide or poly(ethylene glycol) offer a high degree of tunability, but rapidly dissipate forces over short distances (< 25 μm) away from the cell due to their continuum and affine mechanical behavior (*30, 31*). Consequently, we lack an understanding of how the distinct topographical and mechanical properties of fibrous ECMs regulate long-range force transmission and resulting MIC between cells.

Our lab has previously developed synthetic matrices of electrospun dextran-based hydrogel fibers with user-defined architecture and mechanical properties (*32–36*). Here, we combined this biomaterial system with a microfabrication-based cell-patterning method to investigate EC force-mediated matrix displacements and MIC as a function of matrix stiffness. Using this approach, we found that soft, deformable fibrous matrices support long-range matrix deformations and MIC between pairs of ECs. Specifically, we observed that force transmission across fibers spanning neighboring cells was required to support directed protrusions, migration, and the formation of cell-cell contact between ECs up to 200 μm apart. We also identified critical roles for intracellular calcium (Ca^2+^) signaling, focal adhesion kinase (FAK) signaling, and mechanosensitive ion channels during the generation and response to mechanical signals transmitted through the ECM. Lastly, we extended these observations to 3D fibrous settings by examining MIC during vascular network assembly in fibrin hydrogels.

## RESULTS

### Cell-generated matrix deformations and tension support enhanced cell spreading and multicellular cluster formation

Vasculogenic assembly is one context where a deeper understanding of long-range mechanical communication could prove invaluable (*37*). Occurring naturally during embryonic development and adult neovascularization, this process involves assembly of individual ECs into an interconnected network of capillary-like structures and requires cellular communication and coordination over length-scales larger than a cell (*38, 39*). If better understood, control over this process presents a promising approach to engineer microvasculature to support parenchymal cells, a major challenge in the field of tissue engineering and regenerative medicine (*40*). We previously found that cell force-mediated matrix reorganization portends EC network assembly, implicating matrix stiffness in MIC (Fig. 1a,b) (*33*). To more closely examine interactions between ECs as a function of fibrous matrix stiffness, matrices of electrospun DexMA fibers were suspended over an array of microfabricated wells (*32*). Prior to cell seeding, the bulk stiffness of matrices was modulated via photocrosslinking and fibers were functionalized with an RGD-containing peptide to facilitate integrin-dependent cell attachment (Fig. 1a).

**Figure 1.**
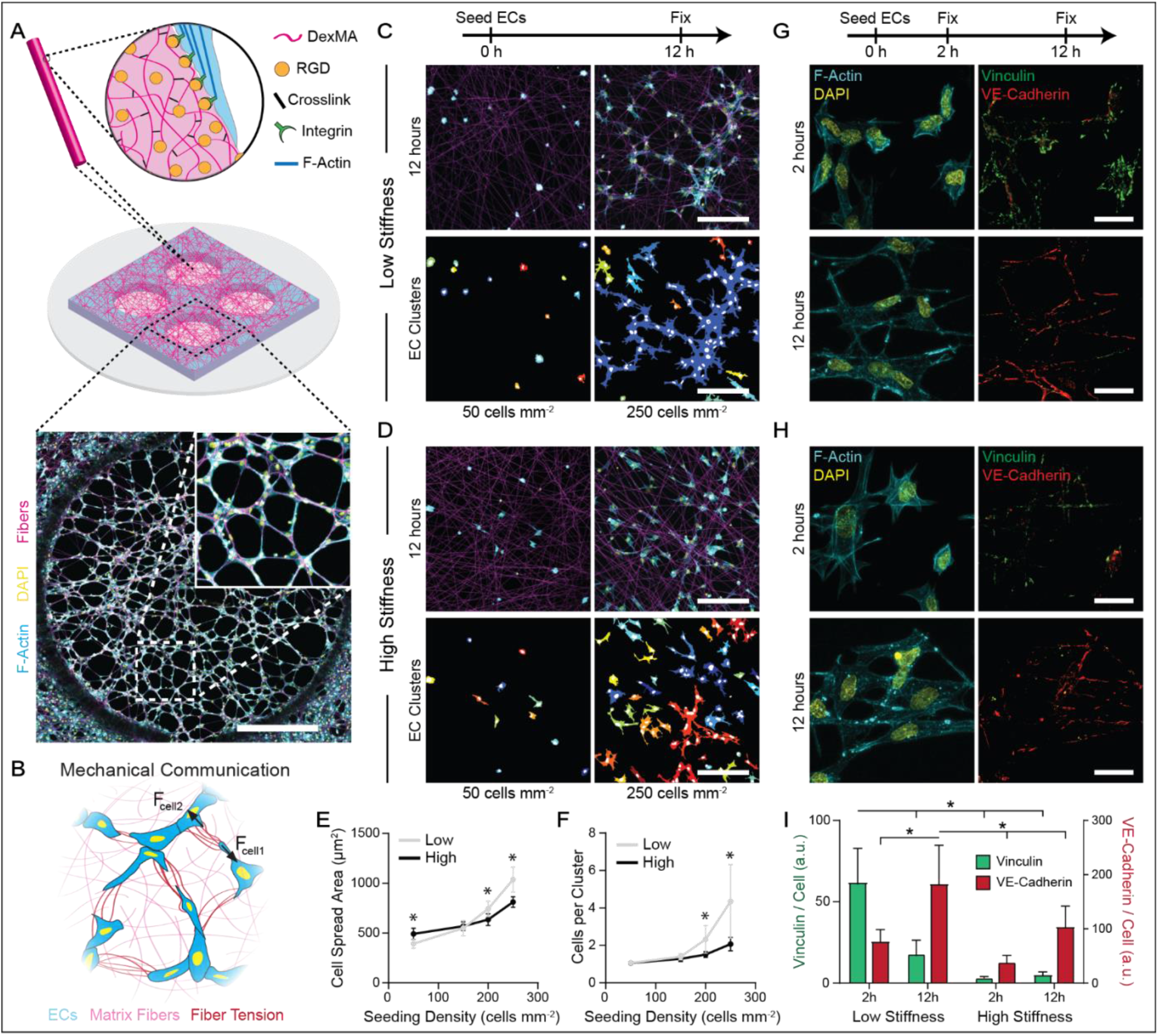
Cell-generated tensile forces and resulting matrix deformations correspond to increased cell spreading and the formation of multicellular clusters. (A) Schematic of microfabricated PDMS multi-well substrate containing an array of wells, each supporting an isolated suspended matrix of DexMA fibers functionalized with RGD to facilitate cell adhesion (scale bar, 1 mm). (B) Schematic of hypothesis that matrix fibers enable MIC underlying EC network formation. (C-D) Confocal fluorescent images of phalloidin-stained ECs (cyan), nuclei (yellow), and rhodamine-labeled DexMA fibers (magenta) with respective color-coded maps of contiguous actin clusters at low (50 cells mm^-2^) and high (250 cells mm^-2^) seeding density in (C) low stiffness, cell-deformable matrices and (D) high stiffness, non-deformable matrices (scale bar, 200 μm). (E) Quantification of cell spread area and (F) average number of ECs per contiguous actin cluster as a function of seeding density and matrix stiffness (n = 12 fields of view). (G-H) Confocal fluorescent images of phalloidin-stained ECs (cyan), nuclei (yellow), vinculin (green), and VE-cadherin (red) at 2 and 12 hours after seeding in (G) low stiffness and (H) high stiffness matrices (scale bar, 25 μm). (I) Quantification of total vinculin and VE-cadherin fluorescent intensity normalized to cell density as a function of time and matrix stiffness (n = 6 fields of view). All data presented as mean ± SD; asterisk denotes significance with P < 0.05.

To test our hypothesis that cell forces and resulting matrix deformations mediate MIC between ECs, we first seeded cells over a range of densities on low stiffness matrices that were deformable (E = 0.724 kPa) or high stiffness matrices that were non-deformable (E = 19.7 kPa) under cellular traction forces (Fig. S1). As seeding density roughly corresponds to the average distance between neighboring cells, we hypothesized the effect of MIC would manifest as cell-density dependent differences in cell spreading on deformable vs. non-deformable matrices. At low seeding densities (50 cells mm^-2^), isolated cells remained largely unspread independent of matrix stiffness, although a moderately higher spread area was noted in non-deformable matrices (Fig. 1c-e). However, increasing seeding density resulted in more marked increases in cell spread area in deformable matrices as compared to non-deformable matrices, with significant differences noted at 200 and 250 cells mm^-2^, indicating a synergistic effect between matrix stiffness and seeding density on cell spreading (Fig. 1e). Along with this increase in spread area, larger interconnected multicellular clusters formed on cell-deformable matrices (Fig. 1f). Although this experiment did not control for paracrine effects which likely are operative, these results suggest that the spacing between cells and sensing of cell force-mediated matrix displacements influences cell spreading and the formation of interconnected, multicellular clusters.

We next investigated cell-ECM and cell-cell adhesion during this process by immunostaining for vinculin (a force-sensitive component of FAs) and VE-cadherin (the direct link between ECs at adherens junctions), respectively (Fig. 1g,h). At an early time point when cells are putatively sending and receiving mechanical signals that mediate cell spreading and directional extension (2 h post-seeding), we observed significantly more vinculin-rich FAs in deformable compared to non-deformable matrices, suggesting heightened cell-ECM adhesion and force transmission to the matrix (Fig. 1i). VE-cadherin levels were low independent of matrix stiffness at this time point, likely due to insufficient time for cells to form stable cell-cell junctions (Fig. 1i). At 12 h post-seeding, however, we observed significantly higher VE-cadherin signal in ECs in deformable matrices compared to those in non-deformable matrices (Fig. 1i). Together, these results indicate that cell-deformable fibrous matrices facilitate strong cell-ECM adhesions that presage the formation of robust adherens junctions and further support the involvement of cell-generated forces transmitted between cells through the matrix during intercellular communication.

The above results suggest low stiffness, cell-deformable fibrous ECM promotes FA formation and generation of mechanical signals that underly cellular communication. To investigate the dynamics of EC assembly into multicellular clusters, timelapse imaging of Hoechst-labeled ECs transduced with a LifeAct-GFP F-actin reporter was conducted over the same 12 h timeframe. As in our previous experiments, ECs actively recruited matrix fibers and formed large multicellular clusters in deformable matrices; this phenomenon was not observed in non-deformable matrices (Movie S1). ECs additionally migrated overall faster in deformable matrices compared to in non-deformable matrices (Fig. 2a-c). Analyzing migration speed over discrete intervals of time, we noted the largest discrepancy in speed (1.5-fold) during the first two hours of culture, compared to a 1.25-fold difference throughout the remainder of timelapse imaging (Fig. 2b,c). Interestingly, enhanced migration speeds over the first two hours coincided with a rapid increase in multicellular cluster size (Fig. 2d). We noted instances of directed extension and migration of neighboring cells towards each other to form multicellular clusters in deformable matrices, while migration in non-deformable matrices appeared uncoordinated or random (Fig. 2e, Movie S1). Together, this data suggests that ECs respond to mechanical signals transmitted through cell-deformable ECM by directionally migrating towards one another at higher speeds to more efficiently form multicellular structures.

**Figure 2.**
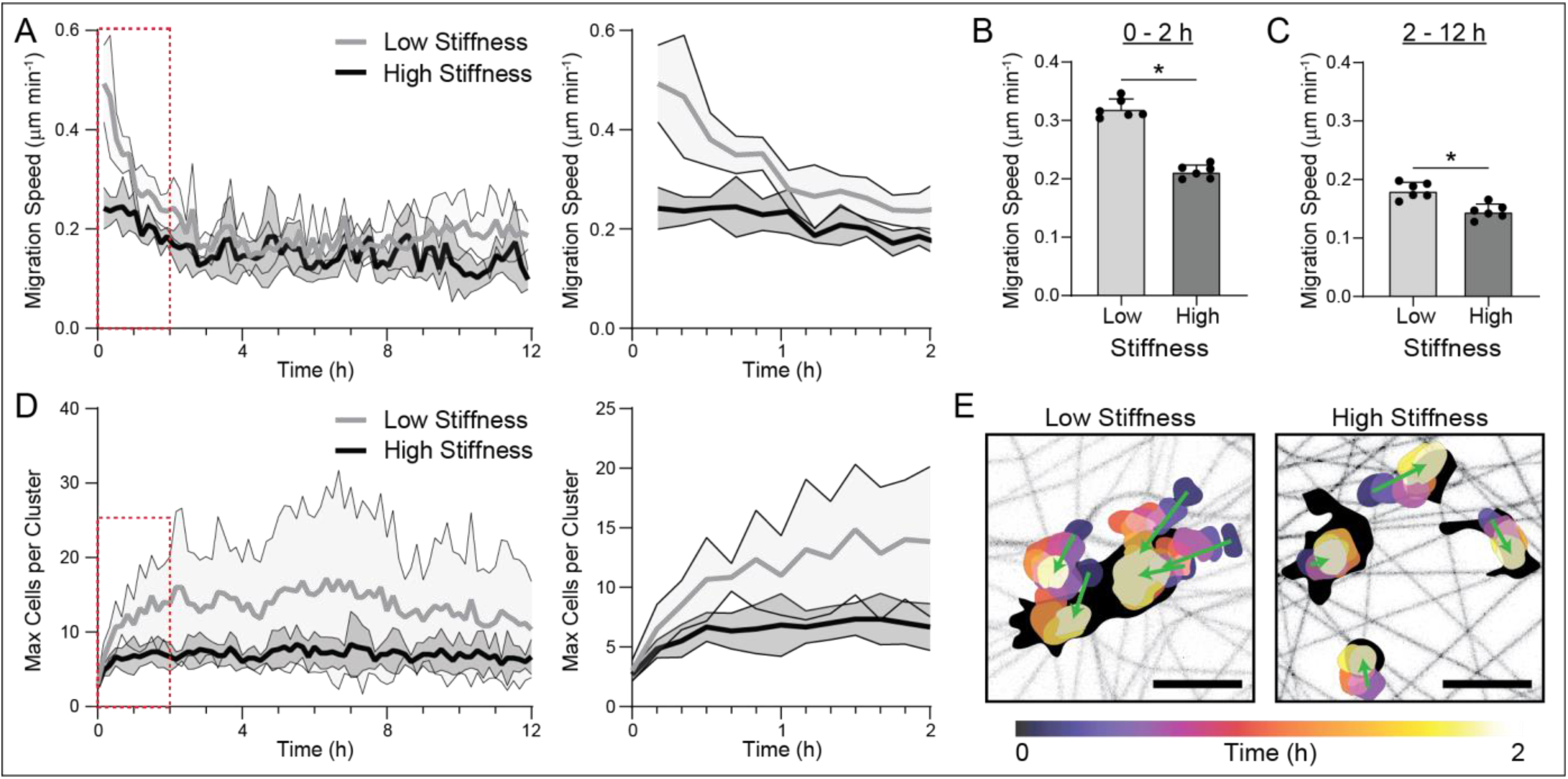
Low stiffness, cell-deformable fibrous matrices increase cell migration speed during multicellular cluster formation. (A) Migration speed of ECs over 12 hours following seeding in either low stiffness, cell-deformable matrices or high stiffness, non-deformable matrices (n = 6 fields of view). (B) Quantification of migration speed during the first 2 hours and (C) remaining 10 hours of culture as a function of matrix stiffness (n = 6 fields of view). (D) Maximum cluster size over 12 hours of culture as a function of matrix stiffness (n = 6 fields of view). (E) Temporally color-coded overlay capturing the motion of nuclei over a 2 hour time course in low and high stiffness matrices. Green arrows represent direction of movement for each individual nuclei with eventual contiguous actin structures demarcated in black (scale bars, 50 μm). All data presented as mean ± SD with superimposed data points; asterisk denotes significance with P < 0.05.

### Micropatterning single ECs reveals matrix stiffness influences cell spreading, FA formation, and matrix deformations

While the previous model provides evidence for MIC in deformable fibrous matrices, the highly dynamic and reciprocal nature of generating, receiving, and responding to mechanical signals within a randomly distributed population of cells is challenging to dissect. Specifically, heterogeneous cell seeding in this setting precludes measuring strain fields of individual ECs, which could provide insight into the generation and transmission of mechanical signals. Thus, we developed a microfabrication-based cell patterning method to precisely pattern individual ECs at the center of a suspended DexMA fiber matrix with defined properties (Fig. 3a, Fig. S2) (*41*). Briefly, ECs were isolated using cell-patterning molds containing an array of 30 μm diameter microwells, approximately the size of a suspended EC following trypsin/EDTA treatment. This cell patterning mold was then aligned with an identically spaced fiber matrix array such that single ECs were accurately and consistently patterned at the center of an individual suspended matrix with a 500 µm diameter (Fig. 3a,b).

**Figure 3:**
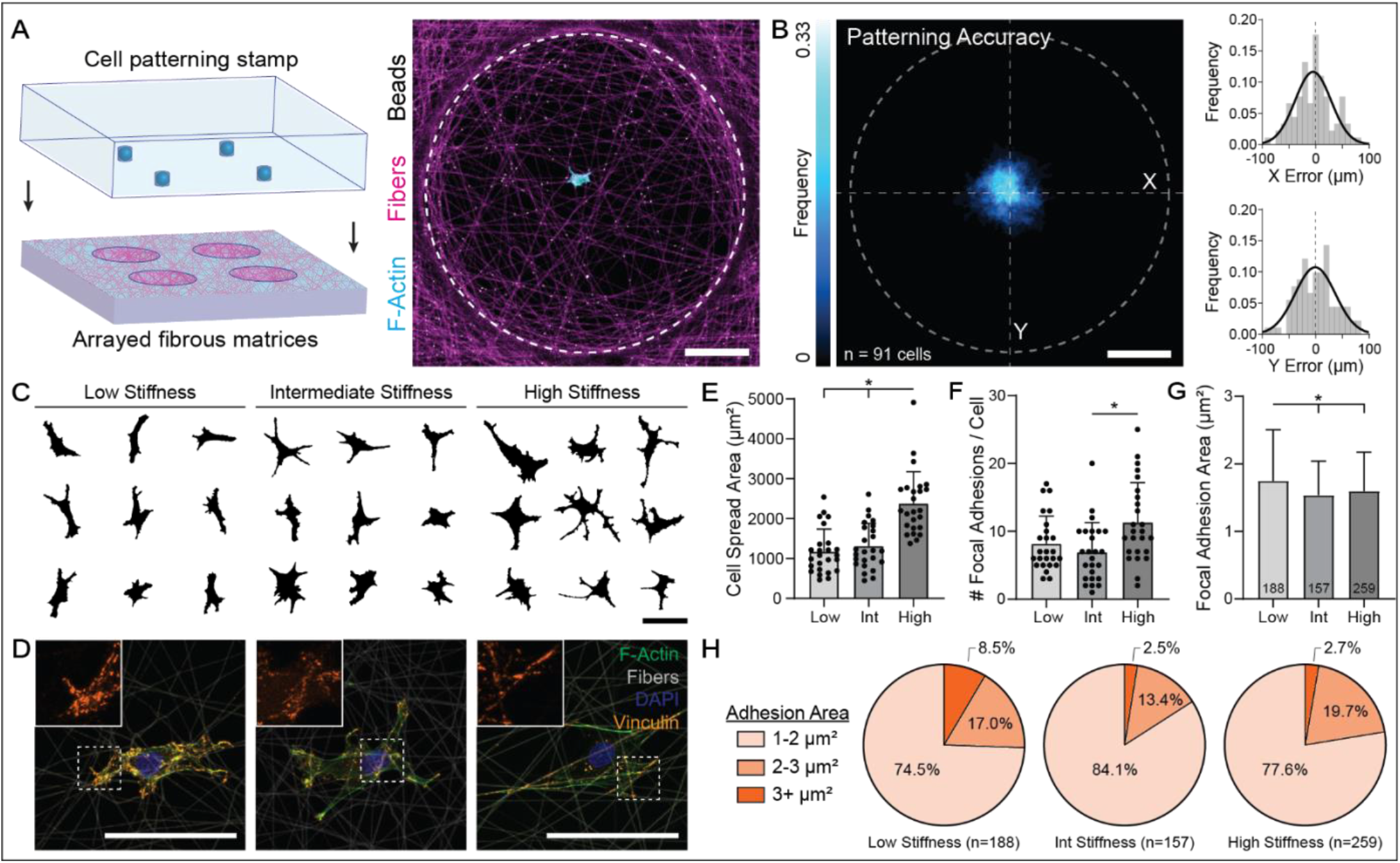
Micropatterning single ECs in individual suspended fibrous matrices reveals cell spreading and FA formation are matrix stiffness-dependent. (A) Schematic depicting microfabrication-based patterning approach to isolate individual ECs at the center of suspended fibrous matrices. Representative confocal fluorescent image of patterned EC (cyan), rhodamine-labeled fibers (magenta), and fluorescent beads embedded in matrix fibers (white) (scale bar, 100 μm). (B) F-actin heat map of patterned ECs with histograms of average patterning error in x-and y-directions (n = 91 cells) (scale bar, 100 μm). (C) Cell outlines of nine representative cells as a function of matrix stiffness (scale bar, 100 μm). (D) Representative immunostained images of FAs in ECs as a function of matrix stiffness; F-actin (green), DexMA fibers (grey), nuclei (blue), and vinculin (orange) (scale bar, 100 μm). (E) Quantification of cell spread area (n = 25 cells), (F) number of FAs per cell (n = 25 cells), and (G) average FA area as a function of matrix stiffness. Data presented as mean ± SD with superimposed data points; asterisk denotes significance with P < 0.05. (H) Distribution of individual FA area as a function of matrix stiffness showing a larger population of 3+ μm^2^ FAs in low stiffness matrices.

Single ECs were patterned in low stiffness/cell-deformable (E = 0.724 kPa), intermediate stiffness (E = 3.15 kPa), or high stiffness/non-deformable (E = 19.7 kPa) DexMA matrices, cultured for 12 hours, and analyzed for cell spreading (F-actin) and FAs (vinculin). With increasing matrix stiffness, we observed a slight increase in cell spread area (Fig. 3c,e) in agreement with previous observations at low EC seeding density (Fig. 1e, 50 cells mm^-2^). Accordingly, we saw a modest increase in the total number of FAs per cell with increasing matrix stiffness (Fig. 3d,f). However, when we analyzed the size of each individual FA, we observed that ECs in soft matrices had significantly higher average FA area despite a lower total number of adhesions (Fig. 3g). This was attributed to a higher proportion of large adhesions (> 3 μm^2^) in low stiffness matrices (8.5%) compared to in intermediate (2.5%) and high (2.7%) (Fig. 3h). Interestingly, the difference in adhesion area as a function of stiffness when ECs were isolated as single cells is modest compared to the difference seen with bulk seeding (Figure 1), suggesting that robust adhesions in low stiffness matrices could be developed as a result of active mechanical signals transmitted by neighboring cells.

While analysis of cell morphology and FAs at fixed timepoints provides useful information, it does not capture the dynamic interplay between cell spreading and ECM deformations. Thus, we next combined our cell/ECM patterning technique with timelapse confocal microscopy (Movie S2). The dynamics of cell spreading and matrix deformations varied with stiffness, specifically in terms of spreading as well as the number and lifetime of protrusions (Fig. 1a,b). In cell-deformable matrices, ECs generally remained unspread for the first four hours of culture during which cells actively recruited matrix fibers beneath the cell body; in contrast, ECs in intermediate stiffness and non-deformable matrices spread immediately (Fig. 4e). In all stiffness conditions, migration was limited and did not significantly vary with matrix stiffness (Fig. 4g). With increasing matrix stiffness, ECs generated more protrusions over the 12-hour timelapse (Fig. 4h, Fig. S3), while the average lifetime of each protrusion decreased (Fig. 4i). As cell protrusions and constituent FAs comprise a critical force-generating apparatus of the cell, these data suggest that cell-deformable matrices not only promote increased FA area (Fig. 1, 3), but also more directional and longer lasting mechanical signals. In addition, we confirmed expected differences in EC force-mediated matrix deformations (Fig. 4c,d,f). In low stiffness matrices, bead displacements were measurable across the entire suspended matrix (up to 250 μm away from the cell centroid) (Fig. 4c,d,l, Fig. S4) and furthermore, temporally correlated with increases in cell spreading (Fig. 4e,f). With increasing matrix stiffness, however, the magnitude and range of displacements diminished, where high stiffness matrices displayed negligible displacements across the entire matrix (Fig. 4k,l).

**Figure 4:**
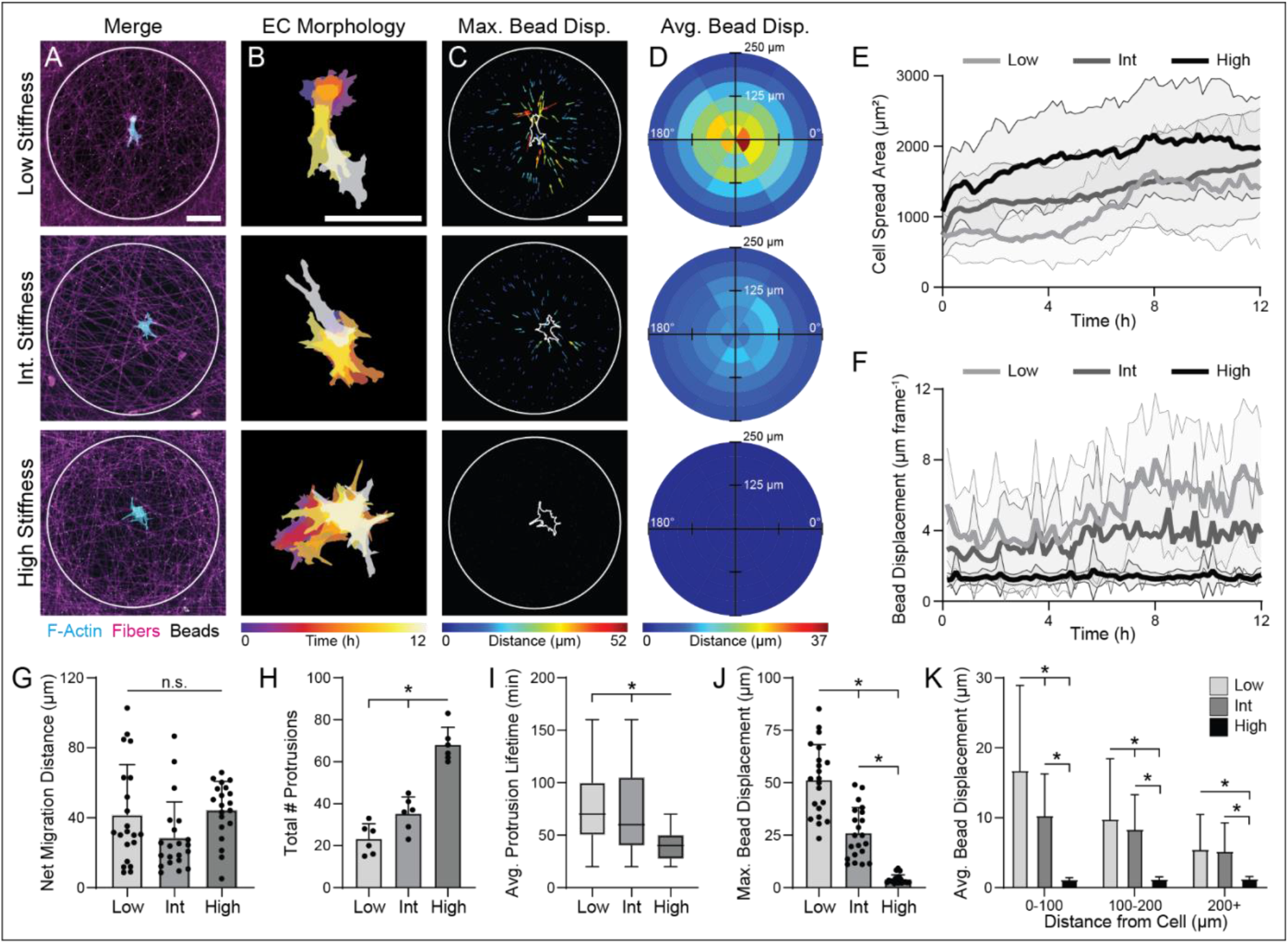
Cell-deformable fibrous matrices support more persistent mechanical signals by promoting fewer, but longer lasting protrusions. (A) Representative confocal fluorescent image of LifeAct-GFP expressing ECs (cyan), rhodamine-labeled DexMA fibers (magenta), and fiber-embedded fluorescent beads (white) (scale bar, 100 μm). (B) Temporally color-coded overlay of EC cell bodies over a 12 hour time course following patterning (scale bar, 100 μm). (C) Size-and color-coded vector plots displaying maximum displacement of each bead over a 12 hour time course (scale bar, 100 μm). (D) Binned average bead displacements for all ECs aligned along their long axis (0°) with color-coded magnitudes (n > 20 cells). (E) Cell spread area and (F) fluorescent bead displacement over a 12 hour time course as a function of matrix stiffness (n > 20 cells). (G) Net migration distance (n > 20 cells), (H) total number of protrusions (n = 6 cells), (I) average protrusion lifetime (n = 6 cells, n = 30 protrusions), and (J) maximum bead displacement as a function of matrix stiffness (n > 20 cells). (K) Binned average bead displacements as a function of starting distance from the cell centroid (n > 20 cells, n > 627 beads). All data presented as mean ± SD with superimposed data points; asterisk denotes significance with P < 0.05.

In addition to stiffness, we also investigated the effect of matrix fiber density by altering the duration of electrospun fiber collection while maintaining a constant degree of crosslinking (equivalent to the lowest stiffness condition above) (Fig. S1). Similar to increasing the stiffness of a fixed density of fibers, increasing the density of low stiffness fibers decreased the magnitude and range of displacements, although to a lesser degree (Fig. S5, Movie S3). Taken together, this data indicates that low stiffness, low density fibrous matrices support long-range matrix deformations and prime ECs for directed force generation by promoting the formation of larger FAs and fewer but longer-lived protrusions.

### Propagation of mechanical signals between neighboring cells promotes directed migration and formation of cell-cell connections

Single cell patterning studies indicate that low stiffness matrices prompt ECs to generate larger, more directional matrix displacements, but the generation of a mechanical signal is only an initial step during MIC. Thus, we next investigated how ECs receive and respond to cell-generated force transmission through fibrous matrices. To do so, we adapted our cell/ECM patterning technique to pattern two cells at a defined distance of 200 μm away from each other (Fig. 5a), as single cell studies demonstrated that ECs in low stiffness matrices generate strain fields that propagate a distance of 200 μm away from the cell (Fig. 4). Rigid boundaries of the microfabricated well were redesigned to ensure even spacing between cell pairs and the fixed boundary at the well edge (Fig. 5a). Combining this approach with timelapse confocal microscopy, we next aimed to determine if and how the stiffness of fibrous matrices regulated cell-cell interactions.

**Figure 5:**
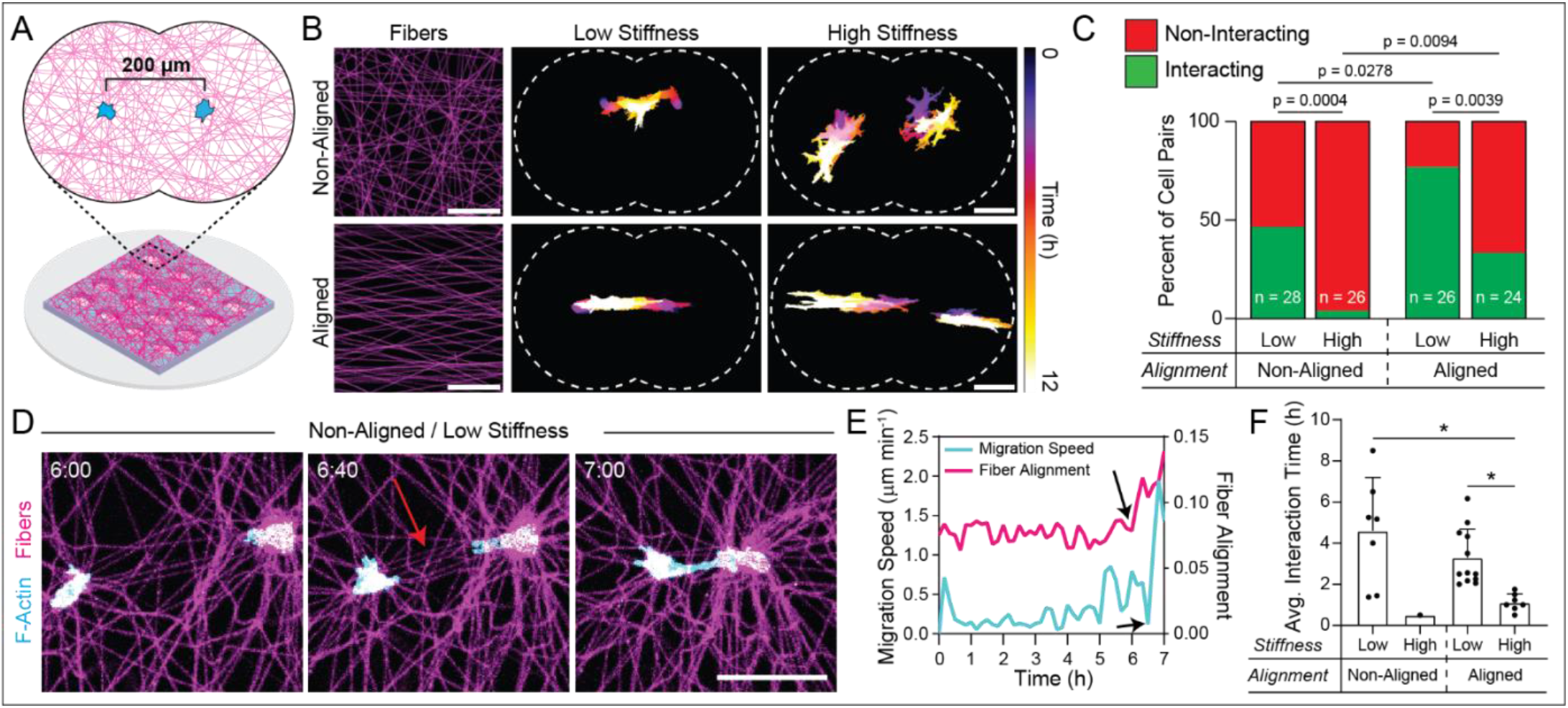
Force transmission through aligned fibers spanning neighboring cells promotes directed migration and the formation of cell-cell connections. (A) Schematic of cell pair patterning in which two ECs are patterned 200 μm apart on a suspended DexMA fibrous matrix. The fiber well is designed such that the distance between the two cells is equal to the distance between cells and well edges. (B) Representative image of initial matrix fiber morphology in non-aligned and aligned conditions. Temporally color-coded overlays capturing EC morphology and migration over a 12 hour time course after cell attachment as a function of matrix alignment and stiffness (scale bars, 100 μm). (C) Quantification of the percent of interacting and non-interacting cells over a 12 hour time course, with interacting cells defined as cell-cell contact at any point during the 12 hour timelapse. P-values determined by Fisher’s exact test. (D) Representative cell pair interaction between ECs patterned in a non-aligned, low stiffness, cell-deformable matrix (scale bar, 100 μm) and (E) quantification indicating an increase in fiber alignment between the two cells followed by a rapid increase in migration speed prior to cell making direct contact. (F) Quantification of average interaction time (duration during which direct cell-cell contact was maintained). All data presented as mean ± SD with superimposed data points; asterisk denotes significance with P < 0.05.

We first patterned cell pairs in cell-deformable and non-deformable DexMA matrices and defined cell-cell interaction as two ECs forming direct cell-cell contact at any time over a 12 h period (Movie S4). In deformable matrices, 46.4% of EC pairs (n = 28) exhibited cell-cell interactions, as compared to 3.9% of EC pairs in high stiffness matrices (n = 26) (Fig. 5b,c). Closer examination of EC pairs in deformable matrices revealed alignment of fibers spanning cell pairs that appeared to mediate directed extension, migration, and resulting cell-cell contact (Fig. 5d). The alignment of fibers appeared to result from cell force-mediated matrix reorganization and preceded directional migration of cell pairs towards one another by approximately 20-30 minutes (Fig. 5e, Fig. S6). Aligned fibers spanning cell pairs did not occur in instances where cells did not interact, independent of matrix stiffness, suggesting that cell force-mediated fiber alignment is a prerequisite to mechanical communication (Fig. S6).

Two non-mutually exclusive explanations for a requirement of fiber alignment in cell-cell interactions include: 1) aligned fibers promote contact guidance cues, enabling directional migration of cell pairs towards each other, and/or 2) aligned fibers maximize force transmission between cells, in turn promoting directed extension, migration, and interaction. To ascertain the relative importance of these two scenarios, we patterned cell pairs on pre-aligned matrices of either low or high stiffness (Fig. 5b, Movie S5). If contact guidance alone were sufficient for cell-cell interactions, we would expect no effect of matrix stiffness on the percent of interacting cells. However, 76.9% of EC pairs (n = 26) in cell-deformable aligned matrices exhibited cell-cell interactions, in contrast to 33.3% of EC pairs in stiffer, non-deformable aligned matrices (n = 24) (Fig. 5b,c). While interactions occurred in all matrix conditions, we noted that the nature of interaction differed between low and high stiffness matrices regardless of matrix alignment. In deformable matrices, ECs maintained cell-cell contact for significantly longer durations compared to in non-deformable matrices (Fig. 5d). Together, these results indicate that force transmission through aligned fibers spanning neighboring cells maximizes the transmission of intercellular mechanical signals to facilitate direct cell-cell contact.

### Patterning EC lines identifies a critical role for coordinated intracellular Ca^2+^ signaling during MIC

These insights could enable the formation and stabilization of organized and stable multicellular structures from larger population of cells in larger tissues with translational potential. To explore this possibility, we further modified our cell/ECM patterning technique to pattern lines of equally spaced cells within a 1 mm² square suspended fiber matrix (Fig. 6a). Cells were patterned approximately 40 μm apart within each line to ensure overlapping cell-generated strain fields in deformable matrices, and two separate lines of cells were placed 500 μm apart to maximize experimental throughput while minimizing potential interactions between lines. Lines of ECs were patterned on cell-deformable and non-deformable matrices to observe how matrix mechanics and MIC regulated the formation and maintenance of organized multicellular structures. ECs patterned in deformable matrices aligned matrix fibers and formed stable structures resembling the original pattern (Fig. 6b,c). ECs patterned in non-deformable matrices, however, appeared to spread and migrate independently, with limited fidelity to their original pattern as quantified by the proportion of nuclei within the original patterned region after 12 hours of culture (Fig. 6b-d). Additionally, VE-cadherin immunostaining to assess the maturation of cell-cell contacts indicated significantly higher VE-cadherin signal in ECs patterned in deformable matrices as compared to non-deformable matrices, in agreement with our previous observations of bulk-seeded matrices (Fig. 6b,e).

**Figure 6:**
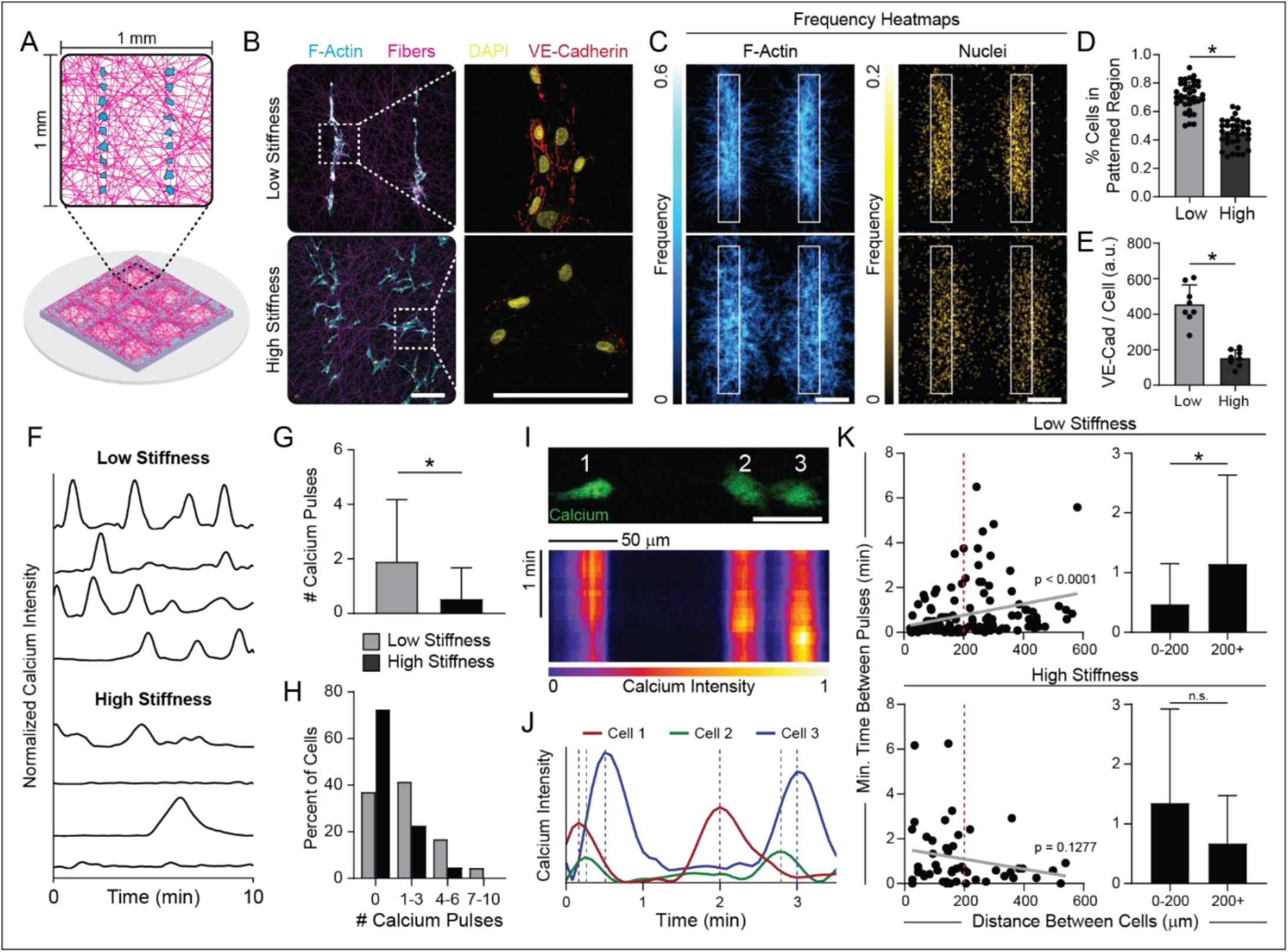
Micropatterned, multicellular lines support a role for coordinated intracellular Ca^2+^ signaling between neighboring cells during MIC. (A) Schematic of EC line patterning consisting of two parallel lines of cells within each suspended fibrous matrices. (B) Representative confocal fluorescent image of phalloidin-stained ECs (cyan), rhodamine-labeled fibers (magenta), nuclei (yellow), and VE-cadherin (red). Dashed boxes indicate locations of higher magnification images of VE-cadherin immunostaining (right column) (scale bars, 200 μm). (C) F-actin (left column) and nuclei (right column) heat maps of EC lines 12 hours after patterning (n = 37 fields of view) (scale bar, 200 μm). (D) Quantification of the percent of nuclei in the original patterned region after 12 hours (n > 35 fields of view). (E) Quantification of total VE-cadherin expression at cell-cell junctions normalized to cell density for each matrix condition (n = 8 fields of view). (F) Normalized Ca^2+^ intensity over a 10 minute time course. (G) Quantification of total number of Ca^2+^ pulses per cell and (H) distribution of cells by number of Ca^2+^ pulses over a 10 minute time frame (n > 89 cells). (I) Representative images of Ca^2+^ signal in ECs within a patterned line with corresponding kymograph and (J) normalized Ca^2+^ intensity displaying a wave of Ca^2+^ fluorescence across the line over time. (K) Quantification of the minimum time between Ca^2+^ pulses as a function of distance between all ECs within patterned lines for low (n = 160 cell pairs) and high stiffness (n = 49 cell pairs) matrix conditions. Grey lines indicate linear correlations with indicated p-values. All data presented as mean ± SD with superimposed data points; asterisk denotes significance with P < 0.05.

Beyond exploring the ability to control patterned multicellular assembly, we also employed this approach to mechanistically explore how ECs communicate with each other during multicellular assembly. Calcium (Ca^2+^) is critical for many cell functions and in particular, Ca^2+^ cytosolic influx has previously been implicated in mechanosensing due to the presence of stretch activated ion channels in many cell types (*42*). Increases in intracellular Ca^2+^ have been shown to trigger a wide variety of cellular processes, including cytoskeletal reorganization underlying cell polarization, protrusion formation, and migration (*43, 44*). Thus, we hypothesized that intracellular Ca^2+^ signaling could play an important role in the EC response to mechanical signals in fibrous matrices.

To examine EC Ca^2+^ signaling, cells were patterned into lines onto suspended fiber matrices, incubated with a Ca^2+^ sensitive dye, and imaged two hours after patterning to visualize intracellular Ca^2+^ flux during the formation of cell-cell connections (Movie S6). Ca^2+^ activity was significantly higher in ECs in cell-deformable matrices (Fig. 6f), with cells exhibiting more frequent Ca^2+^ signal pulses compared to those in non-deformable matrices (Fig. 6g,h). Additionally, we observed instances of temporally sequenced Ca^2+^ pulses between neighboring cells in deformable matrices, where waves of Ca^2+^ fluxed across the line of assembling ECs (Fig. 6i,j). To assess the spatiotemporal regulation of Ca^2+^ signaling within EC lines, we determined the minimum time between Ca^2+^ pulses between two ECs as a function of their separation distance, hypothesizing that MIC would lead to shorter intervals between Ca^2+^ pulses in neighboring cells. Indeed, this correlation was positive and significant in deformable matrices, in stark contrast to non-deformable matrices (Fig. 6k). Interestingly, neighboring ECs within 200 μm from each other (the same distance that interacting cell pairs were patterned in Figure 5) displayed coordinated Ca^2+^ signaling as evidenced by a decreased time interval between Ca^2+^ pulses (Fig. 6l). Neighboring ECs that were further than 200 μm apart, however, had longer average time between peaks with greater variation, indicating a decrease in coordinated Ca^2+^ signaling as a function of distance between cells (Fig. 6l). Together, this data indicates that synchronized Ca^2+^ influx underlies MIC between ECs.

### Focal adhesion kinase signaling and mechanosensitive ion channels are required for MIC during 3D vascular network formation

Given the implication of FAs and Ca^2+^ signaling in MIC (Fig. 1,3), we next inhibited focal adhesion kinase (FAK) as well as the mechanosensitive ion channels (MSICs) transient receptor potential vanilloid 4 (TRPV4) and Piezo1 to examine their roles in Ca^2+^ signaling and the formation of multicellular clusters in deformable fibrous matrices. FAK is an essential non-receptor tyrosine kinase that transduces mechanical signals at FAs to intracellular biochemical signals that coordinate cell behavior (*45*). TRPV4 and Piezo1 are MSICs that open in response to membrane stretch and regulate intracellular Ca^2+^ signaling in ECs (*46, 47*). For patterned cell lines in cell-deformable matrices, inhibition of FAK with PF228 (10 μM) (*48*) led to the largest decrease in intracellular Ca^2+^ signaling, with only 9.5% of all cells exhibiting any Ca^2+^ pulse over a 10 minute time course (Fig. 7a,d,e, Movie S7). Inhibition of TRPV4 with GSK205 (10 μM) (*49*) and Piezo1 with GsMTx-4 (5 μM) (*50*) also led to a decrease in Ca^2+^ signaling compared to ECs in deformable matrices, at comparable levels to ECs in non-deformable matrices (Fig. 7b-e, Movie S7). Additionally, inhibition of FAK, TRPV4, or Piezo1 each led to a decrease in spatially coordinated Ca^2+^ signaling and did not diminish cell force-mediated matrix deformations (Fig. S7,S8). After confirming a role for FAK and MSICs in Ca^2+^ signaling, we seeded ECs (250 cells mm^-2^) in the presence of inhibitors in a 2 mm diameter suspended fiber matrix as in Figure 1 (Fig. 7f). Inhibition of FAK, TRPV4, or Piezo1 led to a decrease in average cell spread area as well as the average number of ECs per cluster after 12 hours of culture in deformable matrices, all exhibiting similar behavior to ECs seeded in high stiffness matrices (Fig. 7g-i). These results indicate that FAK signaling and MSIC activity play an important role in regulating MIC between ECs.

**Figure 7:**
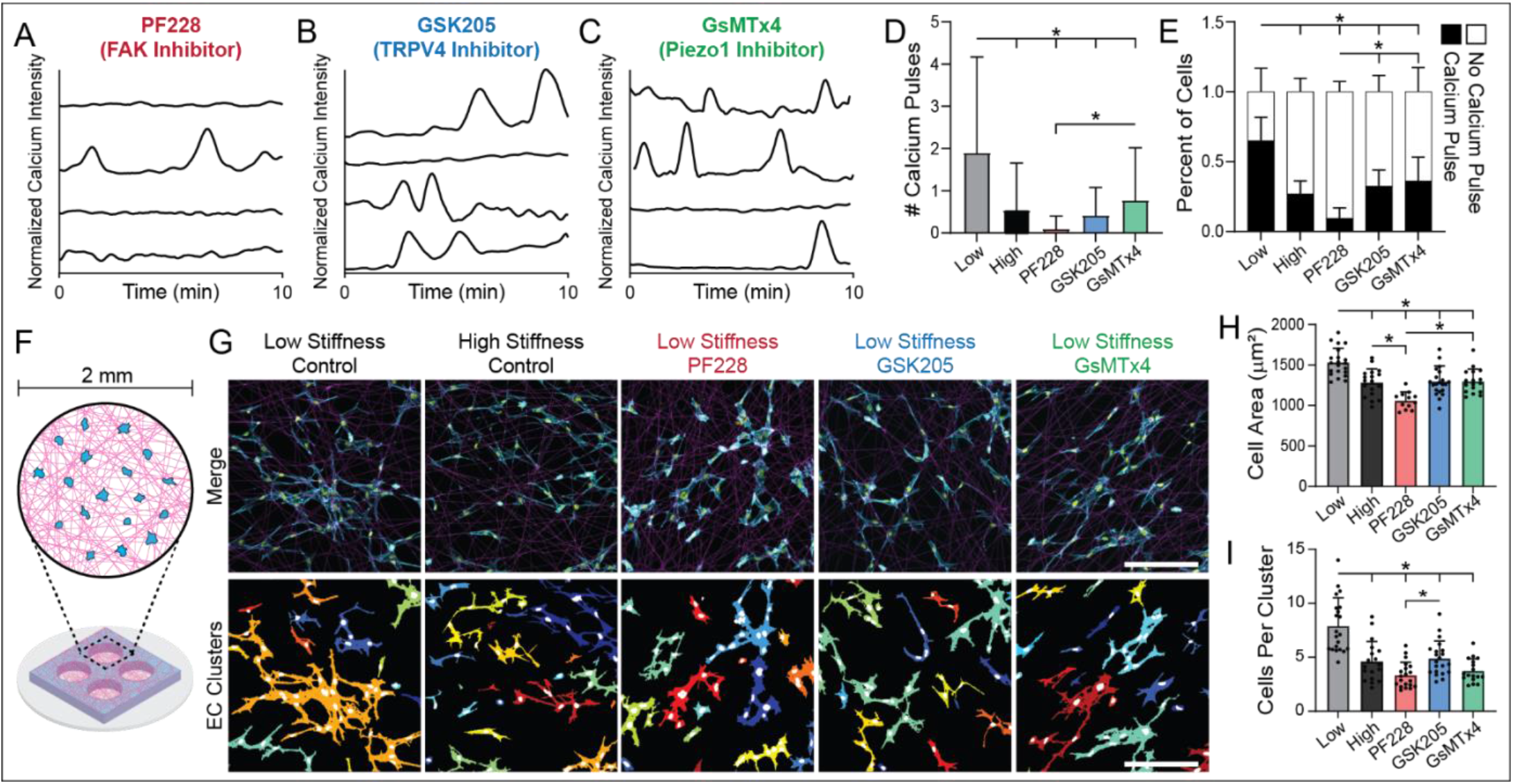
Inhibition of FAK signaling and MSIC activity reduce Ca^2+^ signaling and MIC between ECs. (A-C) Normalized Ca^2+^ intensity over a 10 minute time course of patterned EC lines in low stiffness, cell-deformable matrices treated with (A) PF228 (FAK inhibitor), (B) GSK205 (TRPV4 inhibitor) and (C) GsMTx4 (Piezo1 inhibitor). (D) Quantification of total number of Ca^2+^ pulses per cell and (E) percent of cells with at least one Ca^2+^ pulse over a 10 minute time frame (n > 84 cells). (F) Schematic of EC bulk seeding in 2 mm diameter circular suspended fibrous matrices. (G) Confocal fluorescent images of phalloidin-stained ECs (cyan), nuclei (yellow), and rhodamine-labeled fibers (magenta) with respective color-coded maps of contiguous actin clusters as a function of matrix stiffness and presence of inhibitors (scale bars, 200 μm). (H) Quantification of average cell spread area and (I) average cells per contiguous actin cluster (n > 17 fields of view). All data presented as mean ± SD with superimposed data points; asterisk denotes significance with P < 0.05.

These 2.5D synthetic fibrous matrices provide a controllable setting that allowed us to investigate how biophysical properties of ECM regulate cell behavior and MIC between ECs. However, controlling MIC *in vitro* to engineer functional microvascular networks for tissue engineering applications requires the control of EC network formation in 3D. Using 2.5D suspended fibrous matrices, we identified that ECs utilize mechanical signaling to communicate their location to neighboring cells and found that this process was dependent on FAK signaling and MSIC activity. Whether these findings hold true in a 3D fibrous settings, however, is unclear as such matrices introduce added complexity from the 3D distribution of adhesions and matrix displacements as well as the dependence of cell spreading and protrusions on matrix degradation.

To test this, we examined MIC during vascular network formation in 3D fibrin hydrogels, a commonly used fibrous biomaterial platform for studying vasculogenic assembly (Fig. 8a) (*51*). ECs (4 million cells mL^-1^) were seeded in both 2.5 and 5.0 mg mL^-1^ fibrin hydrogels to model low and high stiffness/density matrix settings, respectively, and cultured for three days. In 2.5 mg mL^-1^ hydrogels, ECs formed long, interconnected networks suggestive of functional microvasculature. ECs in higher density 5.0 mg mL^-1^ hydrogels, however, displayed reduced network formation with limited cell-cell connectivity (Fig. 8b-d, Movie S8). We additionally inhibited FAK and TRPV4 in 2.5 mg mL^-1^ hydrogels, which also led to a marked decrease in average vessel length and interconnected cluster size compared to control (Fig. 8b-d, Movie S8). These results support our findings in synthetic fibrous matrices and imply a critical role for MIC during 3D vascular network formation mediated by FAK signaling and TRPV4 activity.

**Figure 8.**
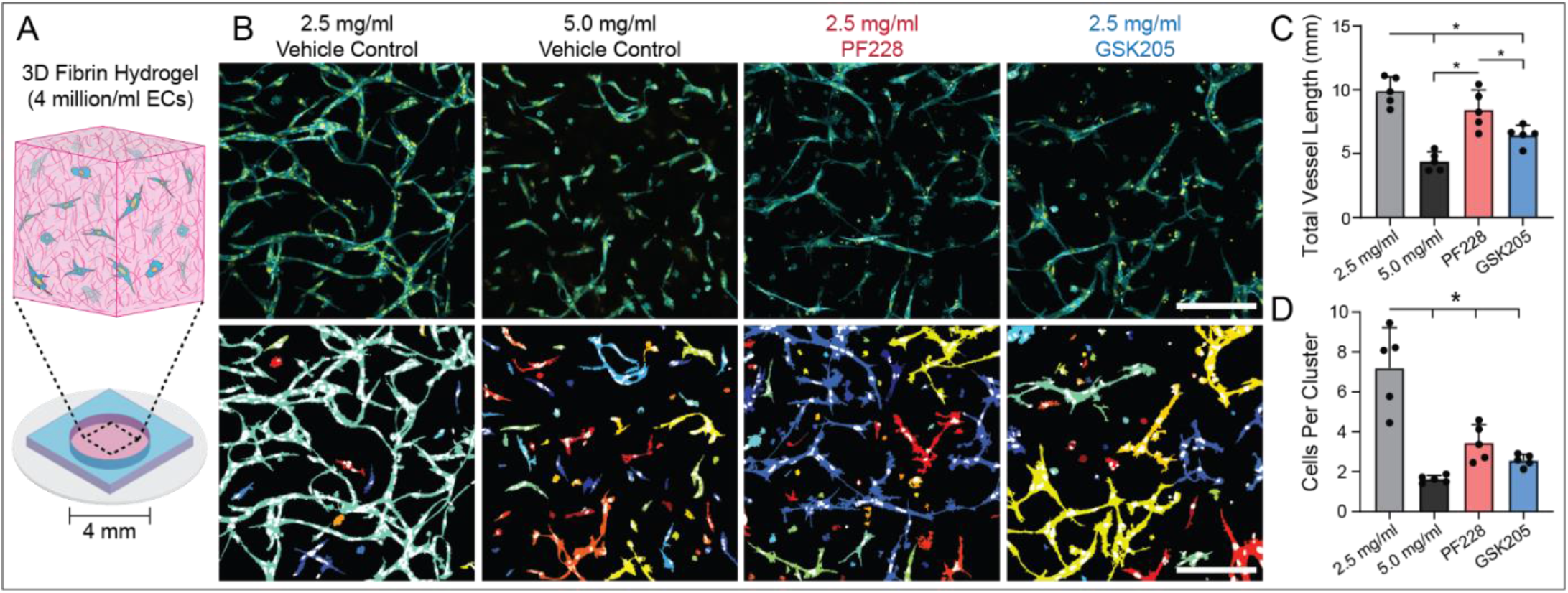
Matrix stiffness/density, FAK signaling, and TRPV4 activity regulate 3D vascular network formation in fibrin hydrogels. (A) Schematic of 3D vascular network formation assay in fibrin hydrogels. (B) Representative confocal maximum intensity projections (100 μm z-stack) of phalloidin-stained ECs (cyan) and nuclei (yellow) with respective color-coded maps of contiguous actin clusters as a function of fibrinogen concentration and presence of inhibitors (scale bars, 200 μm). (C) Quantification of total vessel length and (D) average number of cells in each 3D contiguous actin clusters (n = 5 fields of view). All data presented as mean ± SD with superimposed data points; asterisk denotes significance with P < 0.05.

## DISCUSSION

MIC involves the generation, transmission, and receipt of cell-generated forces conveyed through the ECM, which we posit to be an important but understudied means of intercellular communication. Here, we utilized synthetic fibrous matrices with tunable mechanics in conjunction with a microfabrication-based cell patterning approach to better understand how physical properties of the matrix modulate MIC between ECs. We first varied cell seeding density (and resulting distance between adhered cells) in low and high stiffness fibrous matrices and found that soft, cell-deformable ECM fibers supported heightened cell-ECM adhesion at early time points that corresponded with enhanced cell spreading, migration, and the eventual formation of multicellular clusters (Fig. 1,2). Cell/ECM patterning experiments with single cells and cell pairs revealed that ECs in cell-deformable matrices favored fewer but longer lasting protrusions, generated longer-range matrix displacements, and enabled directional migration towards neighboring cells to form stable cell-cell connections (Fig. 3-6). We also utilized this method to explore the cellular machinery responsible for MIC, finding a critical role for intracellular Ca^2+^ signaling mediated by FAK, TRPV4, and Piezo1 during the assembly of multicellular clusters in deformable matrices (Fig. 6,7). Lastly, we tested whether our observations of MIC in synthetic fiber matrices could be extrapolated to translatable settings by examining 3D vascular network formation in fibrin hydrogels. Indeed, while interconnected vascular-like networks formed in 2.5 mg mL^-1^ fibrin hydrogels, impairing MIC by increasing fibrinogen density (5.0 mg mL^-1^) or FAK/TRPV4 inhibition abrogated the formation of EC networks in 3D fibrous matrices (Fig. 8).

Fibrous matrices have been theorized to promote long-range matrix displacements in a variety of computational models; these studies implicate a role for fiber alignment, strain stiffening, and ECM fiber microbuckling in propagating cell-generated forces (*19–28*). However, exploring cell-force generation and propagation in fibrous matrices *in vitro* is experimentally challenging due the discrete (non-affine), non-linear, and plastic mechanical behavior of naturally derived fibrous ECMs. Thus, we utilized mechanically tunable synthetic fibrous matrices along with timelapse imaging to quantify matrix deformations and found that low stiffness fibrous matrices support longer-range matrix displacements. Additionally, our analysis of cell morphology and cell-ECM adhesion suggest that deformable fibrous matrices prime cells for enhanced and directed force generation and transmission. In experiments with single ECs patterned in deformable matrices, cells exhibited increased FA size and decreased number of protrusions compared to ECs in stiff, non-deformable matrices, despite encountering the same initial ligand density and matrix topography. This may be explained by differences in mechanical resistance provided by the matrix during cell spreading in each matrix condition. In high stiffness matrices, all matrix fibers can provide sufficient resistance required for FA assembly and associated protrusion formation, allowing cells to isotropically extend in an apparently random fashion. In contrast, ECs in low stiffness matrices rapidly recruit matrix fibers soon after initial adhesion, thus generating tension in fibers as a function of their connection to a local rigid boundary (in this case, the edge of the microfabricated well). This likely results in greater heterogeneity of mechanical resistance, leading to preferential generation of FAs and protrusions along particular fibers that provide greater mechanical resistance resulting in directed extension and force propagation. Future work could combine our cell/ECM patterning approach with intramolecular or polymeric tension sensors to better understand the relationship between FA forces and protrusive activity (*52*).

Beyond influencing cell-ECM adhesion, matrix deformations from cell traction forces in low stiffness matrices also led to the formation of structural and mechanical anisotropy via local fiber alignment. When cell pairs were patterned in deformable, randomly oriented fibrous matrices, we observed many instances of fiber alignment between neighboring cells that preceded directional extension, migration, and the formation of cell-cell contact. Several possible explanations exist. Fiber alignment could provide a topographical cue that guides directed migration between cells via contact guidance (*53*). Additionally, and non-exclusively, aligned fibers spanning cells could maximize the transmission of tensile forces between cells and MIC. Our results suggest active force transmission across aligned fibers enhances MIC, as high stiffness aligned matrices that do not deform under cell forces led to fewer cell-cell interactions compared to softer matrices with the same topography (Fig. 5). In recent work, Pakshir et al. observed that contracting myofibroblasts generate large deformation fields in collagen matrices that provide mechanical signals to macrophages (*17*). However, it was also observed that cell force-mediated fiber alignment from myofibroblasts was not required to guide macrophage migration. These conflicting observations may arise from distinct cell types or different ECM settings, as reconstituted collagen hydrogels have relatively short fibrils with nanometer-scale diameters compared to longer electrospun DexMA fibers with larger diameters used in these studies (*33*). Taken together, however, these results confirm that various cell types are able to generate and respond to dynamic mechanical signals in fibrous matrices.

While evidence for MIC has been observed in a variety of settings, the cellular machinery required for cells to send, receive, and respond to mechanical signals has not been established. Here, we identified critical roles for FAK signaling and MSICs in MIC. Inhibition of FAK via PF228 significantly decreased Ca^2+^ signaling and multicellular assembly in matrices permissive to MIC (Fig. 7,8). While FAK inhibition has not been shown to influence FA maturation or traction force magnitudes, it does play an important role in adhesion dynamics and cell motility (*54–56*). Plotnikov et al. investigated FA traction dynamics in mouse embryonic fibroblasts and found FAK activity was required for traction force fluctuations from FAs (*56*). This could in part explain why FAK inhibition hampered Ca^2+^ fluxes in soft matrices, as dynamic traction forces transmitted through the ECM may be critical to MIC (*17*). Furthermore, cells likely sense dynamic mechanical signals from the ECM via membrane stretch resulting in the opening of MSICs. We identified a role for both Piezo1 and TRPV4, two MSICs that are expressed in ECs and are known to be important during the EC response to shear stress (*46, 47*). In addition to regulating EC response to shear stress, though, Thodeti et al. showed that TRPV4 mediates EC reorientation under cyclic mechanical stretch (*46*). Specifically, uniaxial cyclic strain of flexible substrates seeded with ECs led to cellular realignment perpendicular to the axis of applied strain in a TRPV4 dependent manner. The authors posit that ECs preferentially align perpendicular to the direction of applied strain due to differences in membrane strain, TRPV4 activation, and resulting cytoskeletal activity parallel vs. perpendicular to the direction of stretch. This could also explain alignment of cells and directional protrusions in response to cell-generated forces during MIC, as the side of a cell nearest to a force-generating neighboring cell will experience higher levels of membrane stretch. Direct measurements of membrane tension and ion channel state could provide more information on the molecular mechanisms underlying MIC.

MIC is likely critical to tissue development given the ubiquity of cell-generated forces and dynamic changes to the ECM in the growing embryo. The previous notion that cells divide, migrate, and assemble in a static ECM during embryonic development has been widely discredited (*57, 58*). In contrast, collective motion, spatial coordination, and multi-scale ECM deformations are essential for the morphogenetic transitions required for organogenesis (*57, 59, 60*). For example, Zepp et al. recently generated a single-cell RNA-sequencing atlas of the developing mouse lung and identified a critical role for secondary crest myofibroblasts, a highly contractile cell that physically remodels the alveolus and is important for guiding the assembly of alveolar networks (*61*). Additionally, cell forces have been implicated in the folding of mesenchymal tissue during the formation of 3D structures such as gut villi (*62*). Furthermore, timelapse recordings of avian embryonic vasculogenesis demonstrate EC self-assembly into an archetypal polygonal network, termed the primary vascular plexus, prior to the onset of circulation (*63–65*). ECM motion and deformations were noted concurrent to primary vascular plexus formation, potentially indicating a role for MIC between single EC progenitors during this key developmental step.

A complete understanding of developmental vasculogenesis would inform the design of biomaterials that facilitate the self-assembly of functional vascular networks *in vitro* for tissue engineering and regenerative medicine applications. Our studies indicate matrices that permit cellular force transmission via far-ranging matrix deformations promote MIC that is required for the assembly and stabilization of microvascular structures. Interestingly, vasculogenic assembly is readily achieved in collagen and fibrin hydrogels – two materials where evidence of MIC has been well-documented (*15, 17, 51, 66*). These materials, however, hold limited potential for building engineered tissue constructs due to their rapid resorption *in vivo*. Alternatively, synthetic polymeric hydrogels, such as poly(ethylene) glycol, hyaluronic acid, and dextran, offer controllable and modular design better suited for translational applications (*29*). However, compared to the aforementioned natural and fibrous materials, building 3D vascular networks has proven more challenging in these biomaterial settings. Based on our studies, one explanation for the challenges of assembling vascular networks in 3D synthetic hydrogels may be the dissipation of cell forces short distances away from the cell (*30, 31*). Generally, EC spreading and network formation in these settings requires the addition of a support stromal cell such as dermal fibroblasts, mesenchymal stem cells, or pericytes. The role of these stromal cells has long been considered biochemical in nature via growth factor and matrix secretion or ECM degradation (*67–69*). However, in addition to biochemical support, these contractile cells may also provide mechanical signals that orchestrate EC network formation. Supporting this notion, recent work from Song et al. found that fibroblasts present during 3D EC network formation in fibrin is only necessary during the first few days of culture, after which they can be selectively ablated without long-term negative effects on formed vascular networks (*70*).

Additionally, external mechanical cues applied by actuated or dynamic biomaterials could guide cell extension and migration to form complex multicellular patterns (*71*). Beyond playing an important role during the bottom-up assembly of multicellular vascular structures, our results indicate a role for MIC and force transmission in stabilization and maturation, evidenced here by enhanced VE-cadherin in EC clusters and patterned lines formed in cell-deformable matrices. This should be considered in top-down approaches to vascular tissue engineering, such as 3D bioprinting of ECs (*72*). While bioprinting techniques continually improve in resolution and complexity and now allow for cell-scale patterning, an understanding of how matrix properties influence maintenance of cell-cell adhesions is critical, and biomaterial design should consider not only cellular assembly, but also long-term maintenance of multicellular structures.

## MATERIALS AND METHODS

### Reagents

All reagents were purchased from Sigma-Aldrich and used as received, unless otherwise stated.

### Cell culture and biological reagents

Human umbilical vein endothelial cells (ECs) were cultured in endothelial growth medium (EGM-2; Lonza, Basel, Switzerland) supplemented with 1% penicillin-streptomycin-fungizone (Gibco, Waltham, MA). Cells were cultured at 37°C and 5% CO_2_. ECs were used from passages four to eight in all experiments. For live cell time-lapse imaging, lentiviral transduction of LifeAct-GFP was utilized. For inhibition studies, PF-573228 (10 μM), GSK205 (10 μM; Medchem Express, Monmouth Junction, NJ), and GsMTx-4 (5 μM; Abcam, Cambridge, UK) were supplemented in EGM-2 and refreshed every 24 hours.

### Lentivirus production

pLenti.PGK.LifeAct-GFP.W was a gift from Rusty Lansford (Addgene plasmid #51010). To generate lentivirus, plasmids were co-transfected with pCMV-VSVG (a gift from Bob Weinberg, Addgene plasmid #8454), pMDLg/pRRE, and pRSV-REV (gifts from Didier Trono, Addgene plasmid #12251 and #12253 (*73, 74*)) in 293T cells using the calcium phosphate precipitation method (*75*). Viral supernatants were collected after 48 h, concentrated with PEG-it^TM^ (System Biosciences, Palo Alto, CA) following the manufacturer’s protocol, filtered through a 0.45 μm filter (ThermoFisher Scinetific Nalgene, Waltham, MA), and stored at −80°C. Viral titer was determined by serial dilution and infection of ECs. Titers yielding maximal expression without cell death or detectable impact on cell proliferation or morphology were selected for studies.

### DexMA synthesis

Dextran (MW 86,000 Da, MP Biomedicals, Santa Ana, CA) was methacrylated by reaction with glycidyl methacrylate as previously described (*76*). Briefly, 20 mg of dextran and 2 mg of 4-dimethylaminopyridine was dissolved in 100 mL of anhydrous dimethylsulfoxide (DMSO) under vigorous stirring (300 rpm) for 12 h. 24.6 mL of glycidyl methacrylate was then added and the reaction mixture was heated to 45°C for 24 h. The solution was cooled at 4°C for 1 h and precipitated into 1 L ice-cold 2-isopropanol. The crude product was recovered by centrifugation, redissolved in milli-Q water, and dialyzed against milli-Q water for 3 d. The final product was lyophilized and stored at −20°C until use. DexMA was characterized by ^1^H-NMR. The degree of functionalization was calculated as the ratio of the averaged methacrylate proton integral (6. 174 ppm and 5.713 ppm in D2O) and the anomeric proton of the glycopyranosyl ring (5.166 ppm and 4.923 ppm). As the signal of the anomeric proton of α-1,3 linkages (5.166 ppm) partially overlaps with other protons, a pre-determined ratio of 4% α-1,3 linkages was assumed and the total anomeric proton integral was calculated solely on the basis of the integral at 4.923 ppm. A methacrylate/dextran repeat unit ratio of 0.787 was determined.

### Fiber matrix fabrication

Suspended DexMA fiber matrices were fabricated through electrospinning and soft lithography as previously described (*32*). DexMA was dissolved at 0.5 g mL^-1^ in a 1:1 mixture of milli-Q water and dimethylformamide with 1% (w/v) Irgacure 2959 photocrosslinker and 0.625 mM methacrylated rhodamine (Polysciences, Inc., Warrington, PA). For matrix displacement studies, 10% (v/v) blue carboxylate-modified FluoSpheres (1.0 μm diameter, 2% w/v) was also added. Electrospinning was completed with a custom set-up consisting of a high-voltage power supply (Gamma High Voltage Research, Ormond Beach, FL), syringe pump (KD Scientific, Holliston, MA), and a grounded copper collecting surface enclosed within an environmental chamber held at room temperature and 30% relative humidity (Terra Universal, Fullerton, CA). Electrospinning of DexMA solution was performed at a flow rate of 0.45 mL h^-1^, voltage of 7.0 kV, and gap distance of 6 cm. Fiber density was varied through modulating electrospinning time and relative humidity. To induce fiber alignment, fibers were electrospun at a voltage of 4.0 kV onto a collecting surface of oppositely charged (−3.0 kV) parallel electrodes at a 25 mm separation distance. After electrospinning, fibers were stabilized by primary crosslinking under ultraviolet (UV) light (100 mW cm^-2^) for 60 s, hydrated in varying concentrations of lithium phenyl-2,4,6-trimethylbenzoylphophinate (LAP; Colorado Photopolymer Solutions, Boulder, CO) photoinitiator solution, and then exposed again to UV light (100 mW cm^-2^) for 20 s. Low, intermediate, and high stiffness networks were crosslinked in 0.02, 0.075, and 1.0 mg mL^-1^ LAP solutions, respectively. Fibers were collected on various poly(dimethylsiloxane) (PDMS; Dow Silicones Corporation, Midland, MI) arrays of wells produced by soft lithography. Silicon wafer masters possessing SU-8 photoresist (Microchem, Westborough, MA) were first 27 fabricated by standard photolithography. Briefly, a layer of SU-8 2075 (110 μm thick) was spin-coated on a 3-inch silicon wafer and patterned into arrays of various shaped wells spaced evenly within 12 x 12 mm squares. These masters were utilized to make PDMS stamps which were silanized with trichloro(1H,1H,2H,2H-perfluorooctyl)silane and used to emboss uncured PDMS onto oxygen plasma-treated coverslips. Resultant fiber-well substrates were methacrylated by vapor-phase silanization of 3-(trimethoxysilyl)propyl methacrylate in a vacuum oven at 60°C for at least 6 h to promote fiber adhesion to PDMS.

### Mechanical testing

To determine the Young’s modulus of suspended DexMA fibrous matrices, microindentation testing with a rigid cylinder was performed on a commercial CellScale Microsquisher (CellScale, Waterloo, Ontario). Briefly, samples were indented to a depth of up to 200 μm at an indentation speed of 2 μm s^-1^, and Young’s modulus was approximated assuming the material behaves as an elastic membrane as previously described (*32*).

### RGD functionalization and seeding on DexMA matrices

DexMA fibers were functionalized with the cell adhesive peptide CGRGDS (RGD; Peptides International, Louisville, KY). An RGD concentration of 2 mM was used for all studies. RGD was coupled to available methacrylates via Michael-type addition. Briefly, the peptide was dissolved in milli-Q water containing HEPES (50 mM), phenol red (10 μg mL^-1^), and 1 M NaOH to adjust the pH to 8.0. 250 μL of this solution was added to each substrate and incubated for 30 min at room temperature. Following RGD functionalization, substrates were rinsed 2x with PBS before cell seeding. For bulk seeding of networks, ECs were trypsinized, resuspended in 1.5% (w/v) methylcellulose supplemented EGM-2 to increase media viscosity, and seeded between 50 and 250 cells mm^-2^.

### Fluorescent staining and microscopy

ECs on DexMA fibers were first fixed in 4% paraformaldehyde for 10 min at room temperature. Alternatively, to extract cytoplasmic vinculin, samples were simultaneously fixed and permeabilized in 2% paraformaldehyde in a microtubule-stabilizing buffer containing 1,4-piperazinediethanesulfonic acid (PIPES, 0.1 M), ethylene glycol-bis(2-aminoethylether)-N,N,N’,N’-tetraacetic acid (EGTA, 1 mM), magnesium sulfate (1 mM), poly(ethylene glycol) (4% w/v), and triton X-100 (1% v/v) for 10 min at room temperature. To stabilize the fibers for processing and long-term storage, DexMA samples were crosslinked in 2 mL LAP solution (1 mg mL^-1^) and exposed to UV light (100 mW cm^-2^) for 30 s. To stain the actin cytoskeleton and nuclei, cells were permeabilized in PBS solution containing triton X-100 (5% v/v), sucrose (10% w/v), and magnesium chloride (0.6% w/v), and simultaneously blocked in 1% (w/v) bovine serum albumin and stained with phalloidin and DAPI. For immunostaining, samples were blocked for 1 h in 1% (w/v) bovine serum albumin and incubated with mouse monoclonal anti-vinculin antibody (1:1000, Sigma #V9264) or mouse monoclonal anti-VE-cadherin antibody (1:1000, Santa Cruz #sc-9989) followed by secondary antibody (1:1000, Life Technologies #A21236) for 1 h each at room temperature with 3x PBS washes in between. Fixed samples were imaged on a Zeiss LSM800 laser scanning confocal microscope. Unless otherwise specified, images are presented as maximum intensity projections. Fluorescent images were processed and quantified via custom Matlab scripts.

### Cell migration analysis

Immediately after seeding, substrates were transferred to a motorized and environmentally controlled stage and imaged using a Zeiss LSM800 laser scanning confocal microscope (Zeiss, Oberkochen, Germany). Prior to imaging, cell nuclei were labeled with Hoechst 33342 (5 μg mL^-1^) for 10 min. F-Actin, DexMA fibers, and Hoechst-labeled nuclei were imaged at 10 min frame intervals over 12 hours. Following raw image export, images were converted to maximum intensity projections, and cell spreading and cluster formation were quantified using a custom Matlab script. Nuclei tracking was completed with TrackMate, a freely available ImageJ plugin (*77*). Nuclei were detected at each time point using a Laplacian of Gaussian (LoG) detector with an estimated particle diameter of 20 μm and threshold of 0.01 with use of a median filter. Single particle tracking was completed using a linear assignment problem tracker with a linking max distance and gap-closing distance of 20 μm and gap-closing max frame gap of 2 frames. Tracks were filtered to only contain nuclei detected through the entire time-lapse, and migration speed for each cell was calculated via custom Matlab scripts.

### Microwell patterning stamp fabrication

To pattern single ECs onto suspended matrices of DexMA fibers, we designed a patterning system inspired by a previously developed microwell-based approach (*41*). Like the fiber-well substrates, microwell patterning stamps were produced by soft lithography. First, silicon wafer masters were fabricated with two steps of photolithography. First, SU-8 2075 (100-200 μm thick) was spin-coated on a silicon wafer and patterned into 12.05 x 12.05 mm elevated squares. This step allowed for the final patterning stamp to be aligned to the square fiber-well substrate (12 x 12 mm) during patterning. Next, SU-8 2025 (35 μm thick) was spin-coated on top of the previous exposed layer and patterned into arrays of micro-posts (30 μm diameter) centered onto each 12.05 mm square. Non-exposed SU-8 was washed off through the developing process and these masters were utilized to make PDMS microwell patterning stamps as described above.

### Cell patterning on DexMA matrices

Microwell patterning stamps were sterilized with 70% ethanol and UVO (Jelight Company Inc., Irvine, CA) for 5 minutes followed by treatment with 0.2% Pluronic F127 to prevent cell adhesion. 500 μL of EC suspension (1 x 10^6^ cells mL^-1^) was seeded on the patterning stamp and the cells were allowed to settle into the microwells for 5 min. The supernatant was subsequently removed and excess, untrapped cells were gently flushed away with 4 rinses of PBS and 1 rinse of EGM-2. The fiber-well substrate containing RGD-functionalized DexMA fiber matrices was carefully inverted on top of the patterning stamp and aligned. This assembly was then inverted to allow single ECs trapped in microwells to settle and adhere to the DexMA fiber matrix for 15 minutes (Fig. S2). The final substrate containing patterned ECs on DexMA matrices was then hydrated with EGM-2 supplemented with HEPES (25 mM) to regulate media pH and minimize hydrolysis-mediated fiber degradation during culture.

### Timelapse fiber displacement microscopy

Immediately after patterning, substrates were transferred to a motorized and environmentally controlled stage and imaged using a Zeiss LSM800 laser scanning confocal microscope. Images of F-actin, DexMA fibers, and fluorescent beads embedded in fibers were acquired every 10 min for 12 h. Images were converted to maximum intensity projections before analysis. Single particle tracking was completed by first aligning image stacks of fluorescent beads, cropping the images to only track beads within fibers of the suspended matrix, and tracked using TrackMate. Beads were detected at each time point using a Laplacian of Gaussian (LoG) detector with an estimated particle diameter of 5 μm and threshold of 1.0 with use of a median filter. Single particle tracking was completed using a linear assignment problem tracker with a linking max distance and gap-closing max distance of 20 μm and gap-closing max frame gap of 2 frames. Tracks were filtered to only contain particles detected throughout the entire time-lapse and then analyzed using custom Matlab scripts. Additionally, cell morphology, migration, and protrusions were analyzed using custom Matlab scripts.

### Calcium Imaging

Ca^2+^ handling analysis was performed by incubating cells for 1 h at 37°C with 5 μM Cal520, AM (AAT Bioquest, Sunnyvale, CA) in EGM-2. Cells were then returned to normal EGM-2 and allowed to equilibrate for 30 minutes. Following equilibration, substrates were transferred to a motorized and environmentally controlled stage and imaged using a Zeiss LSM800 laser scanning confocal microscope. Timelapse movies of Ca^2+^ flux were analyzed with custom Matlab scripts (*78*).

### 3D Vascular Network Formation

HUVECs were encapsulated (4 million mL^-1^) in fibrin precursor solutions containing 2.5 mg mL^-1^ or 5.0 mg mL^-1^ fibrinogen from bovine plasma and 1 U mL^-1^ bovine thrombin. This precursor solution was mixed and 20 μL was transferred into a PDMS mold with 4 mm diameter and incubated at 37°C for 20 min. Samples were then hydrated in EGM-2 supplemented with fetal bovine serum (FBS; 5% v/v), vascular endothelial growth factor (VEGF; 50 ng mL^-1^), and phorbol 12-myristate 13-acetate (PMA; 25 ng mL^-1^), and media was replaced every 24 hours. Quantification of morphological network properties was performed on 100 μm image stacks. Total vessel length was quantified using AngioTool (*79*), and three-dimensional analysis of the number cells per contiguous actin structure was completed using a custom Matlab script.

### Statistical Analysis

Statistical significance was determined by one-way or two-way analysis of variance (ANOVA) with post-hoc analysis (Tukey test) or Student’s t-test where appropriate. Additionally, statistical significance of the proportions of interacting and non-interacting cells was determined by Fisher’s exact test. For all studies, significance was indicated by p < 0.05. Sample size is indicated within corresponding figure legends and all data are presented as mean ± standard deviation.

## ACKNOWLEDGEMENTS

We thank Dr. Brenton Hoffmann for insightful discussions and critical feedback on this manuscript.

## Author Contributions

C.D.D. and B.M.B. designed the experiments. C.D.D. performed the experiments and analyzed the data with the help of S.J.D., W.Y.W., J.L.K., and D.K.P.J. C.D.D. and B.M.B. wrote the manuscript. All authors reviewed the manuscript.

## Data and Materials Availability

All data needed to evaluate the conclusions in the paper and/or the Supplementary Materials. Additional data related to this paper may be requested from the authors.

## Supplemental Information

### SUPPLEMENTAL FIGURES

**Figure S1.**
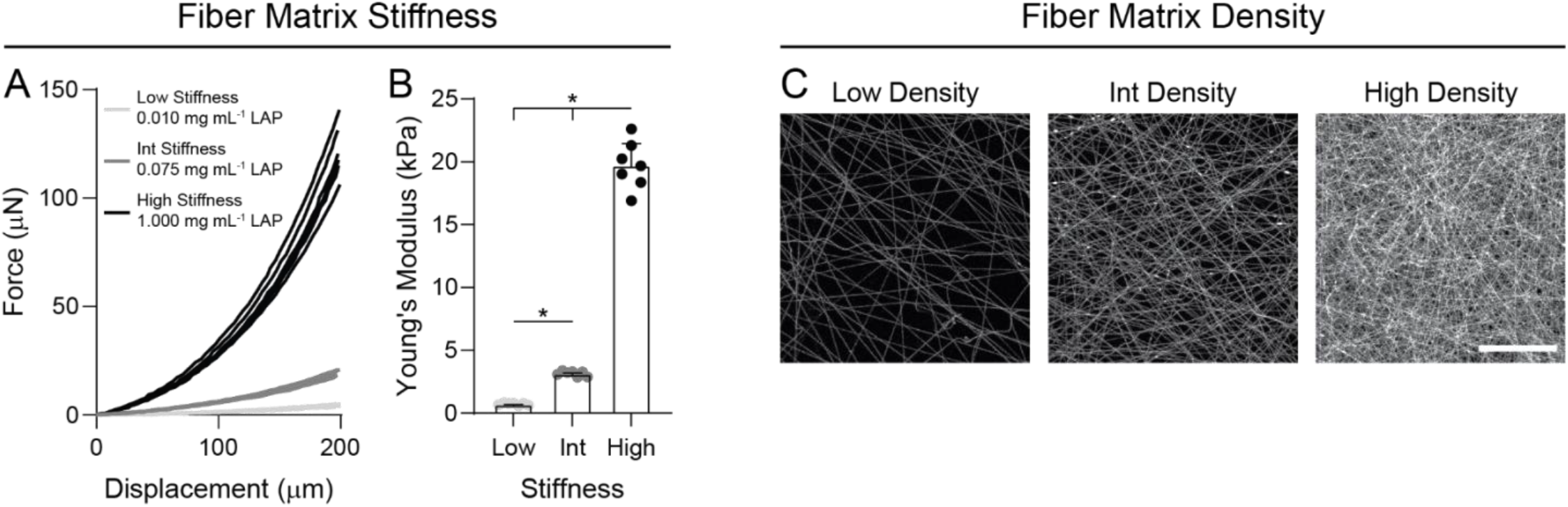
Control over substrate stiffness and fiber density in DexMA fiber networks. (A) Force-indentation response and (B) Young’s modulus of DexMA fibrous matrices as a function LAP photoinitiator concentration (n = 7 matrices/group). (C) Representative fluorescent images of DexMA fibers at low, intermediate, and high fiber density (scale bar, 100 μm).

**Figure S2.**
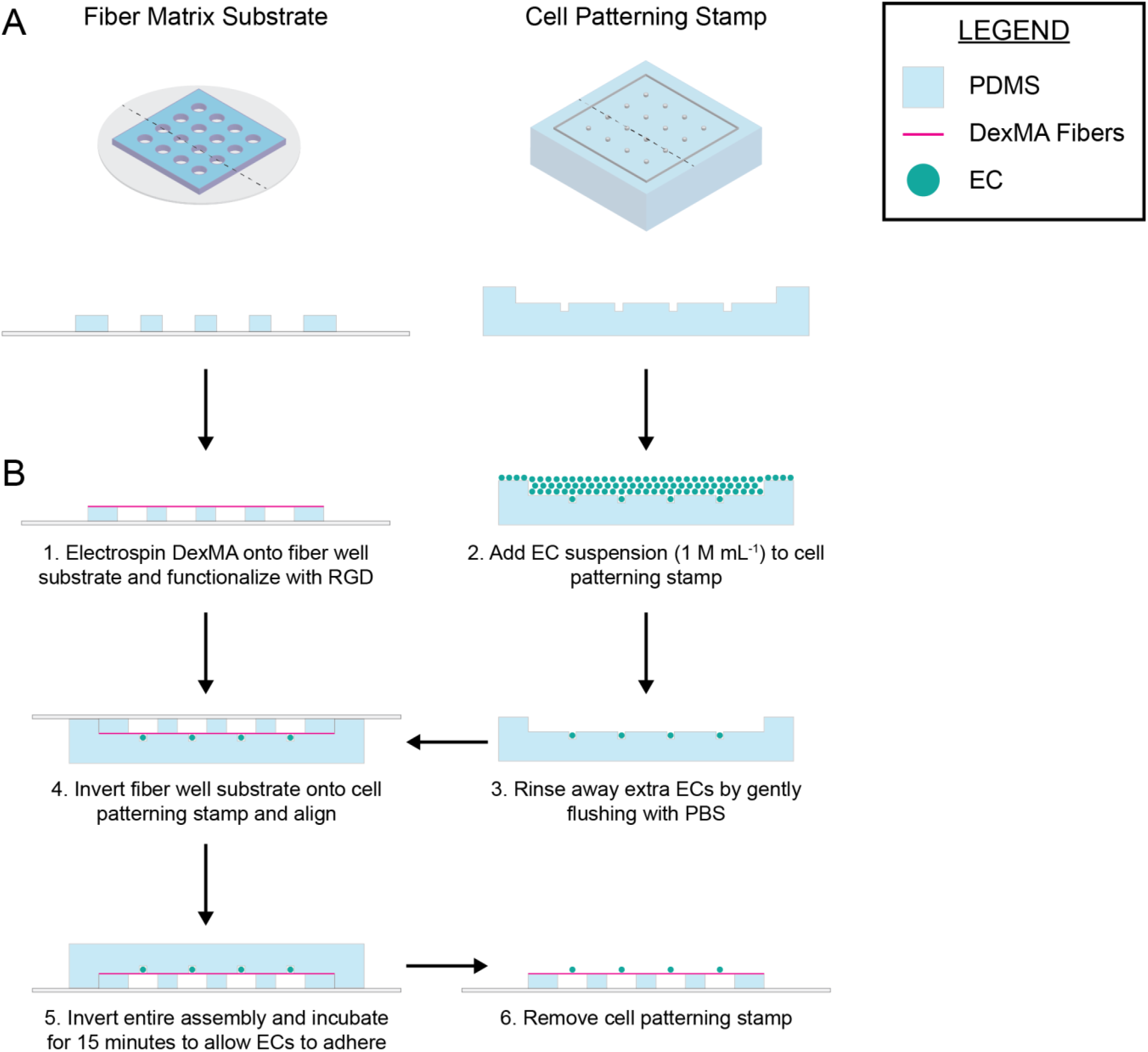
Schematic of endothelial cell micropatterning on suspended DexMA fiber matrices. (A) Isometric and side views of fiber matrix substrate and cell patterning stamp. (B) (1) DexMA fibers are electrospun onto a fiber well substrate, crosslinked, and hydrated. (2) The cell patterning stamp is treated with Pluronic F127 solution to prevent cell adhesion, and a cell suspension is seeded onto the stamp. Cells are allowed to settle for 5 minutes. (3) Excess cells are gently flushed away with 4x PBS rinses. (4) The fiber well substrate is inverted onto the cell patterning stamp and aligned using the raised edges of the patterning stamp under a microscope. (5) The entire assembly is inverted to allow cells to settle onto the suspended DexMA matrices. (6) The patterning stamp is removed and transferred for culture and subsequent imaging.

**Figure S3.**
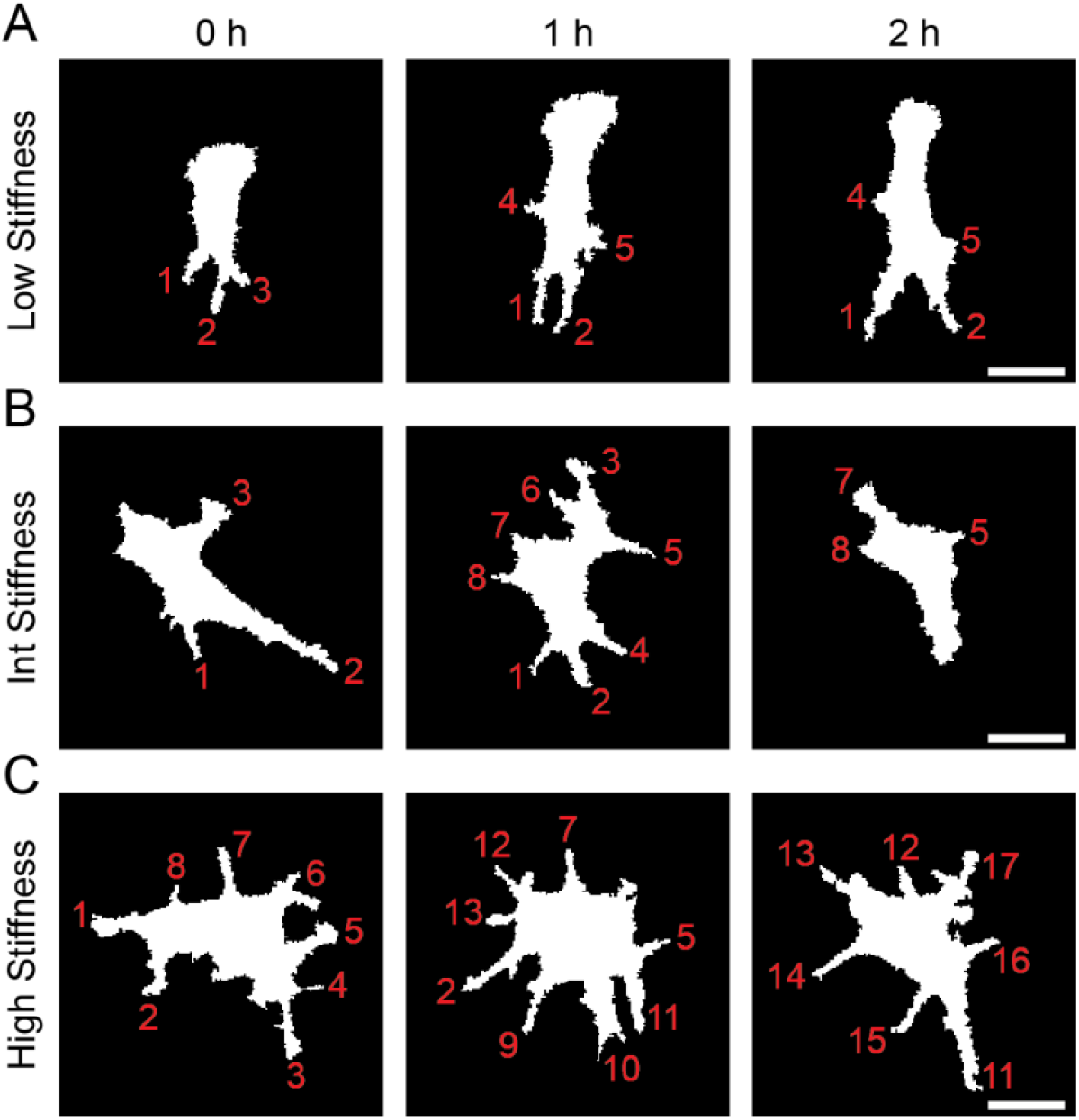
Endothelial cell protrusion analysis. For each frame of the 12 hour timelapse, protrusions were identified and the total number of protrusions as well as lifetime of each individual protrusion was manually determined for cells in (A) low, (B) intermediate, and (C) high stiffness matrices (scale bars, 50 μm).

**Figure S4.**
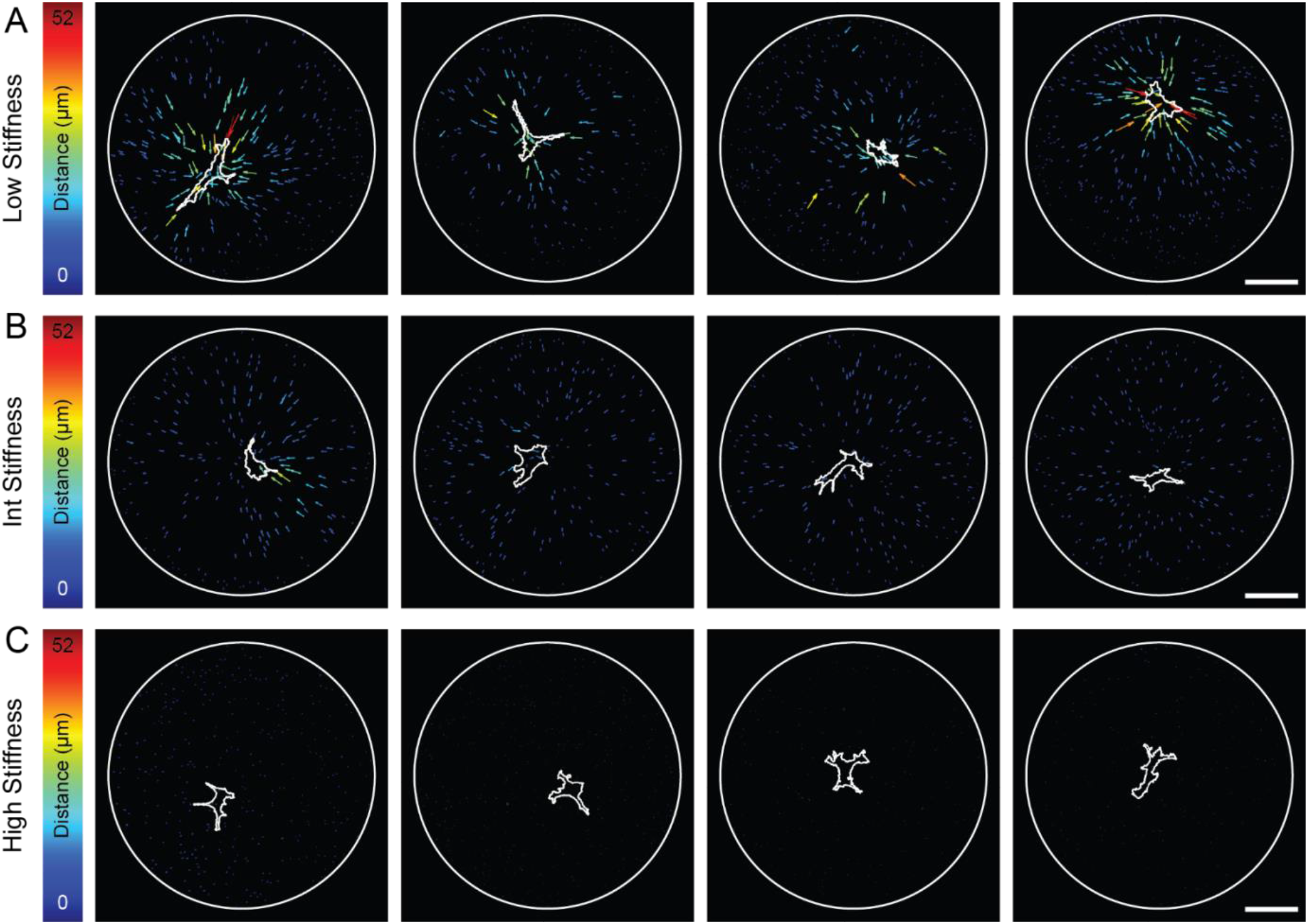
Bead displacement vector plots as a function of matrix stiffness. Representative endothelial cells (cell periphery denoted by white outline) and their respective bead displacements over 12-hours of culture displayed as vectors with magnitude coded by vector size and color in (A) low, (B) intermediate, and (C) high stiffness DexMA matrices (scale bars, 100 μm).

**Figure S5.**
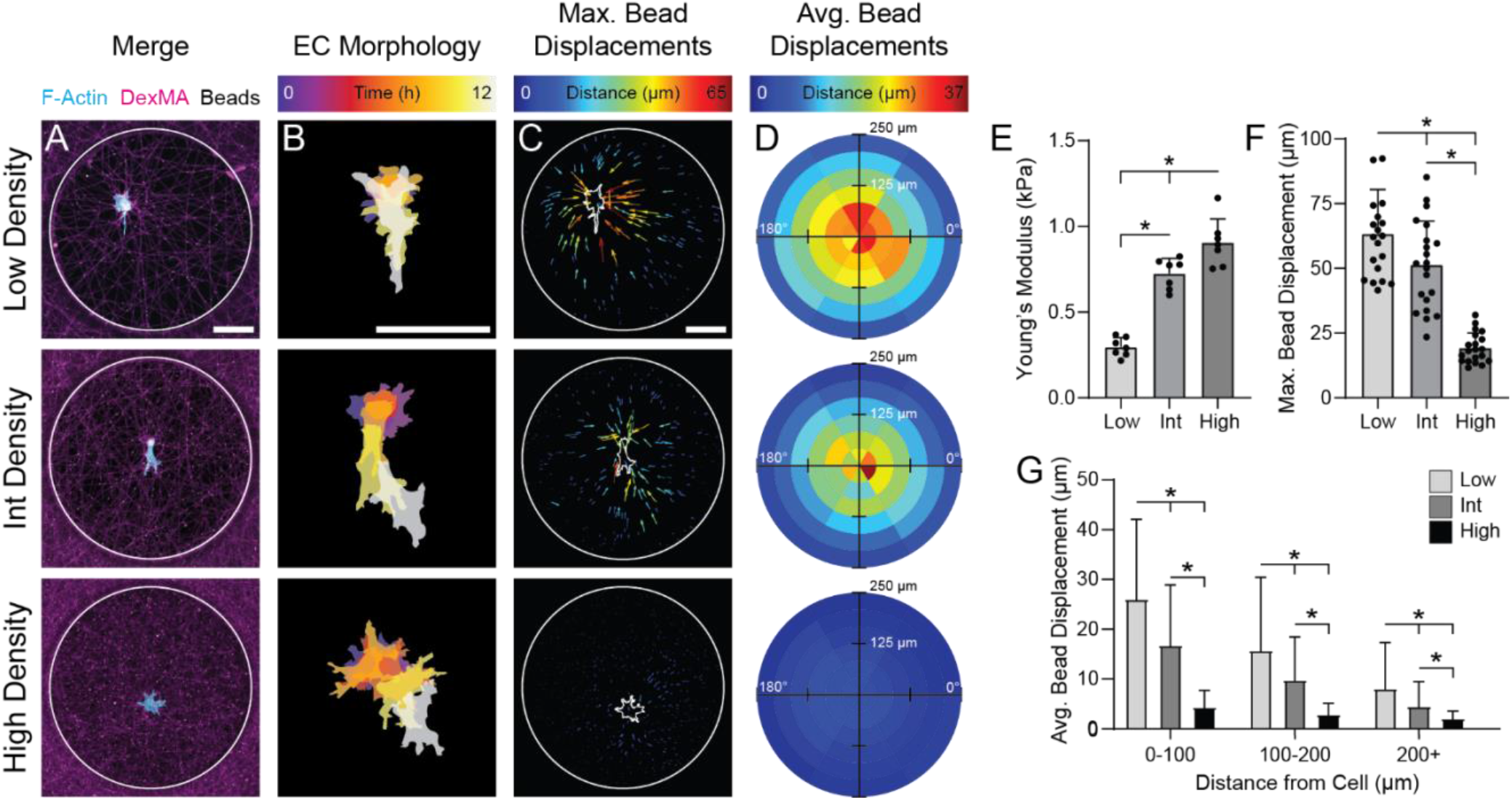
Single endothelial cell morphology and force transmission as a function of fiber density. (A) Representative confocal fluorescent image of phalloidin-stained ECs (cyan), rhodamine-labeled DexMA fibers (magenta), and fiber-embedded fluorescent beads (white) (scale bar, 100 μm). (B) Temporally color-coded overlay of EC cell bodies over a 12 hour time course following initial patterning (scale bar, 100 μm). (C) Maximum displacement of each bead coded by vector length and color over the 12 hour time course (scale bar, 100 μm). (D) Binned average bead displacements color-coded by magnitude for all analyzed ECs with respect to their long axis (0°) (n > 20 cells/group). (E) Young’s modulus of DexMA fiber matrices as a function of matrix density (n = 6). (F) Quantification of maximum bead displacement and (G) binned average bead displacements as a function of starting distance from the cell centroid for low, intermediate, and high fiber densities. All data presented as mean ± SD with superimposed data points; asterisk denotes significance with P < 0.05.

**Figure S6.**
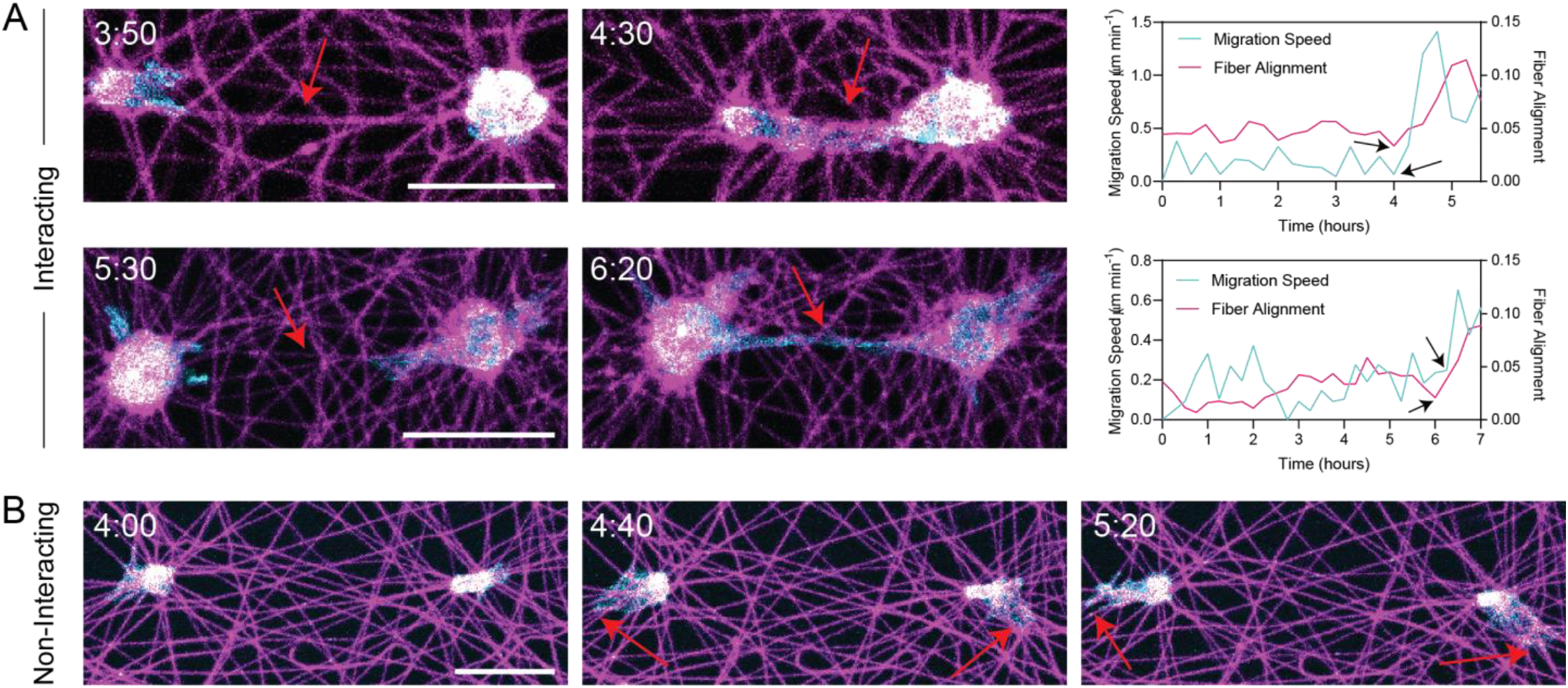
Representative examples of interacting and non-interacting endothelial cell pairs in low stiffness, non-aligned fibrous matrices. (A) Representative confocal fluorescent images of phalloidin-stained ECs (cyan) and rhodamine-labeled fibers (magenta) with quantification of fiber alignment spanning cells and average migration speed of both ECs as a function of time. Red arrows indicate local regions of fiber alignment between neighboring cells (scale bars, 50 μm). (B) Representative confocal fluorescent images of non-interacting ECs. Red arrows indicate cell protrusions extending in the opposite direction of neighboring cell (scale bar, 50 μm).

**Figure S7.**
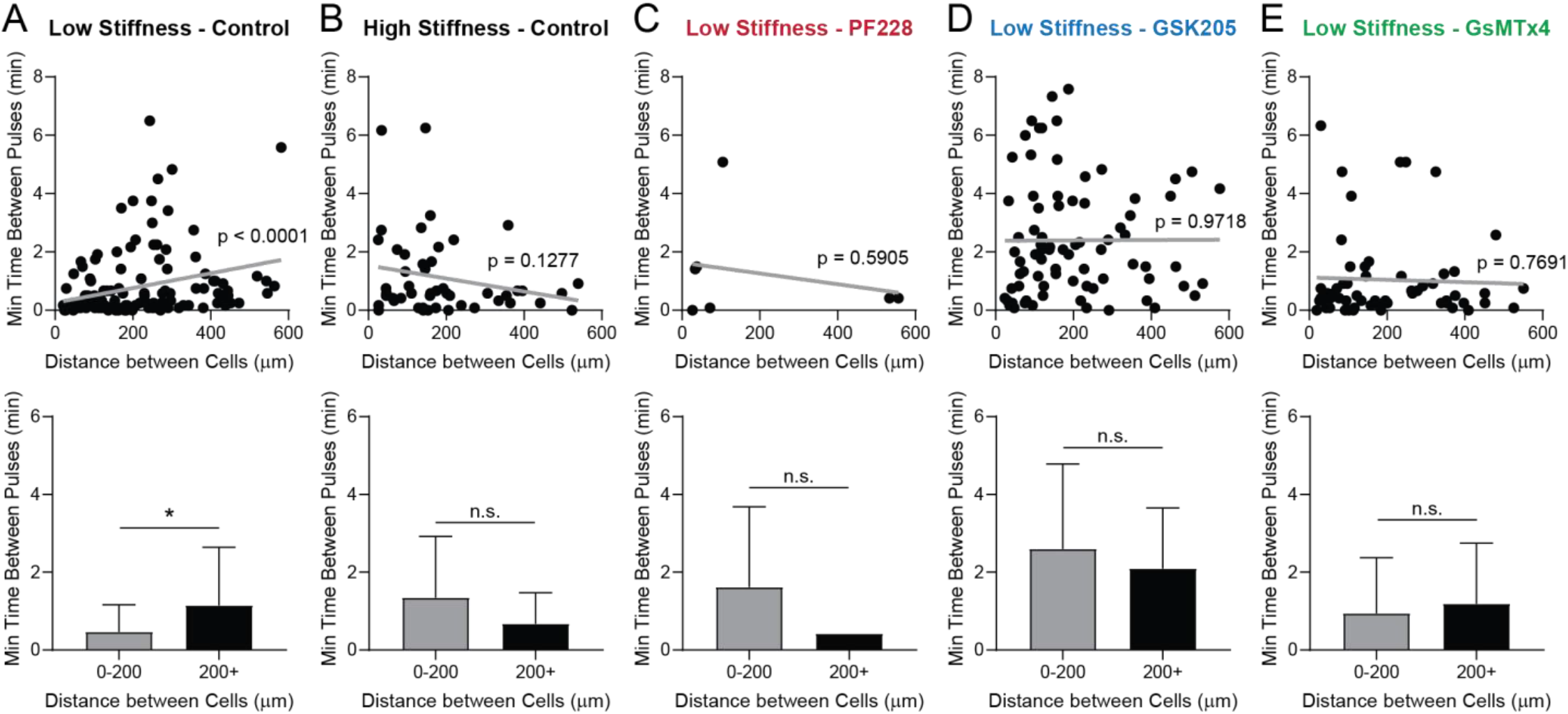
Spatiotemporal Ca^2+^ signaling analysis. Quantification of the minimum time between Ca^2+^ pulses as a function of intercellular distance within EC lines patterned in (A) low stiffness matrices, (B) high stiffness matrices, (C) low stiffness matrices treated with 10 μM PF228, (D) low stiffness matrices treatedd with 10 μM GSK205, and (E) low stiffness matrices treated with 5 μm GsMTx4.

**Figure S8.**
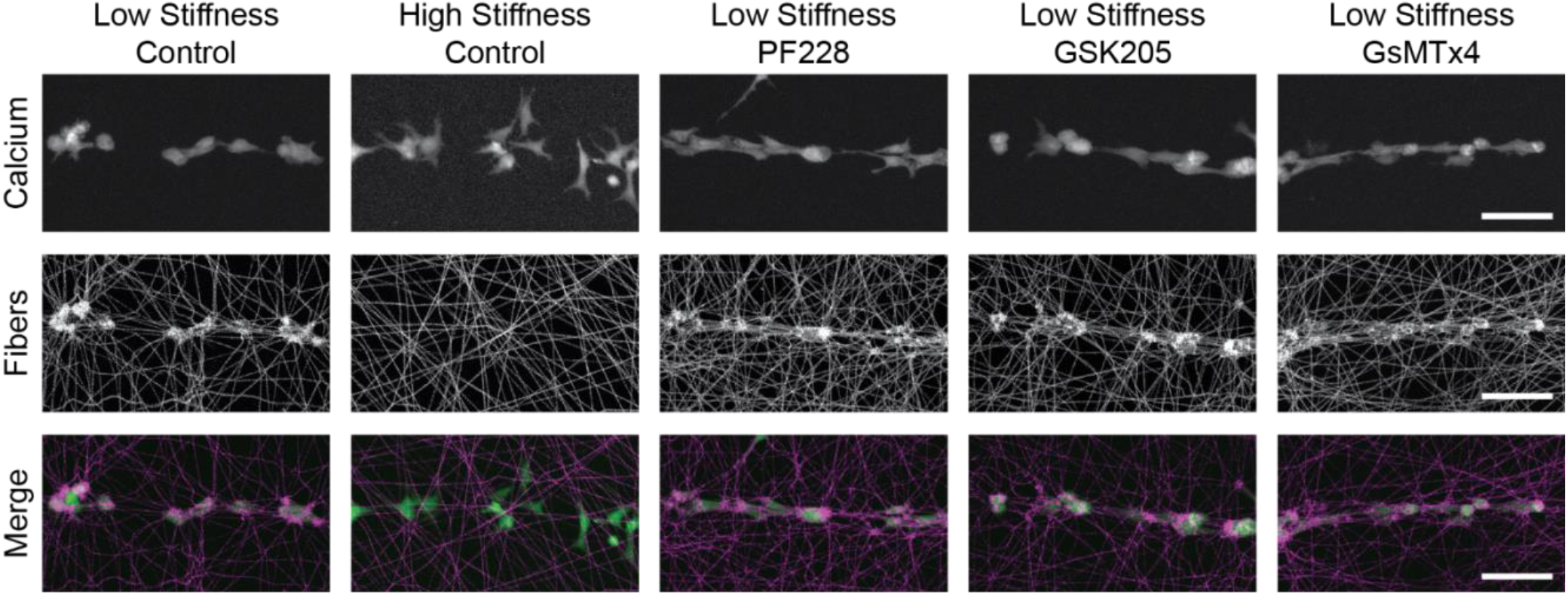
Cell force-mediated matrix deformations following treatment with FAK, TRPV4, and Piezo1 inhibitors. Representative maximum intensity projections of calcium-labeled ECs (green) and rhodamine-labeled fibers (magenta) 2 h after cell line patterning as a function of matrix stiffness and presence of inhibitors confirms that inhibition of FAK (PF228), TRPV4 (GSK205), or Piezo1 (GsMTx4) does not diminish cell force-mediated matrix reorganization (scale bars: 100 μm).

### SUPPLEMENTAL MOVIE CAPTIONS

**Movie S1: EC bulk seeding in fibrous DexMA matrices.** Representative confocal fluorescence 12-hour timelapse movie of lifeAct-GFP expressing ECs (cyan) and rhodamine-labeled DexMA fibers (magenta) seeded at 250 cells mm^-2^ in low stiffness, cell-deformable and high stiffness, non-deformable matrices (Scale bar: 100 μm).

**Movie S2: Single cell patterning in fibrous DexMA matrices with variable matrix stiffness.** Representative confocal fluorescence 12-hour timelapse movie of a single lifeAct-GFP expressing EC (cyan), rhodamine-labeled DexMA fibers (magenta), and embedded fluorescent beads (white) patterned in low stiffness, cell-deformable and high stiffness, non-deformable matrices (Scale bar: 100 μm).

**Movie S3: Single cell patterning in synthetic fibrous DexMA matrices with variable matrix density.** Representative confocal fluorescence 12-hour timelapse movie of a single lifeAct-GFP expressing EC (cyan), rhodamine-labeled DexMA fibers (magenta), and embedded fluorescent beads (white) patterned in low and high density matrices with crosslinking equivalent to the lowest stiffness condition (Scale bar: 100 μm).

**Movie S4: Cell pair patterning in non-aligned fibrous DexMA matrices with variable matrix stiffness.** Representative confocal fluorescence 12-hour timelapse movie of pairs of lifeAct-GFP expressing ECs (cyan) and rhodamine-labeled DexMA fibers (magenta) patterned in low stiffness, cell-deformable and high stiffness, non-deformable non-aligned matrices (Scale bar: 200 μm).

**Movie S5: Cell pair patterning in aligned fibrous DexMA matrices with variable matrix stiffness.** Representative confocal fluorescence 12-hour timelapse movie of pairs of lifeAct-GFP expressing ECs (cyan) and rhodamine-labeled DexMA fibers (magenta) patterned in low stiffness, cell-deformable and high stiffness, non-deformable aligned matrices (Scale bar: 200 μm).

**Movie S6: Calcium transients of ECs in fibrous DexMA matrices with variable matrix stiffness.** Representative confocal fluorescence 10-minute timelapse movie of calcium signaling (green) in ECs patterned into multicellular lines in low stiffness, cell-deformable and high stiffness, non-deformable matrices (Scale bar: 100 μm).

**Movie S7: Calcium transients of ECs in fibrous DexMA matrices in the presence of FAK, TRPV4, and Piezo1 inhibitors.** Representative confocal fluorescence 10-minute timelapse movie of calcium signaling (green) in ECs patterned into multicellular lines in low stiffness, cell-deformable matrices with inhibition of FAK (PF228, 10 μM), TRPV4 (GSK205, 10 μM), and Piezo1 (GsMTx4, 5 μM) (Scale bar: 100 μm).

**Movie S8: Three-dimensional image stack of EC networks in fibrin hydrogels with variable matrix density, FAK inhibition, and TRPV4 inhibition.** Representative confocal fluorescence image stacks (100 μm thick) of phalloidin-stained ECs (cyan) and nuclei (yellow) in 2.5 mg mL^-1^ fibrin, 5.0 mg mL^-1^ fibrin, and 2.5 mg mL^-1^ fibrin with inhibition of FAK (PF228, 10 μM) or TRPV4 (GSK205, 10 μM) (Scale bar: 100 μm).

